# An Lgr5-independent developmental lineage is involved in mouse intestinal regeneration

**DOI:** 10.1101/2024.03.11.584399

**Authors:** Maryam Marefati, Valeria Fernandez-Vallone, Morgane Leprovots, Gabriella Vasile, Frédérick Libert, Anne Lefort, Gilles Dinsart, Achim Weber, Jasna Jetzer, Marie-Isabelle Garcia, Gilbert Vassart

**Affiliations:** Institute of Interdisciplinary Research in Molecular Human Biology (IRIBHM), Université Libre de Bruxelles (ULB), 1070 Brussels, Belgium; Institute of Pathology and Molecular Pathology, University Hospital Zurich, University of Zurich, Switzerland; Institute of Molecular Cancer Research, University of Zurich, Switzerland

## Abstract

Collagenase/dispase treatment of intestinal tissue from adult mice generates cells growing in matrigel as stably replatable cystic spheroids in addition to differentiated organoids. Contrary to classical EDTA-derived organoids, these spheroids display poor intestinal differentiation and are independent of Rspondin/Noggin/EGF for growth. Their transcriptome resembles strikingly that of fetal intestinal spheroids, with downregulation of crypt base columnar cell (CBC) markers (Lgr5, Ascl2, Smoc2, Olfm4). In addition, they display upregulation of inflammatory and mesenchymal genetic programs, together with robust expression of YAP target genes. Lineage tracing, cell-sorting and single cell RNA sequencing experiments demonstrate that adult spheroid-generating cells belong to a hitherto undescribed developmental lineage, independent of Lgr5+ve CBCs, and are involved in regeneration of the epithelium following CBC ablation.

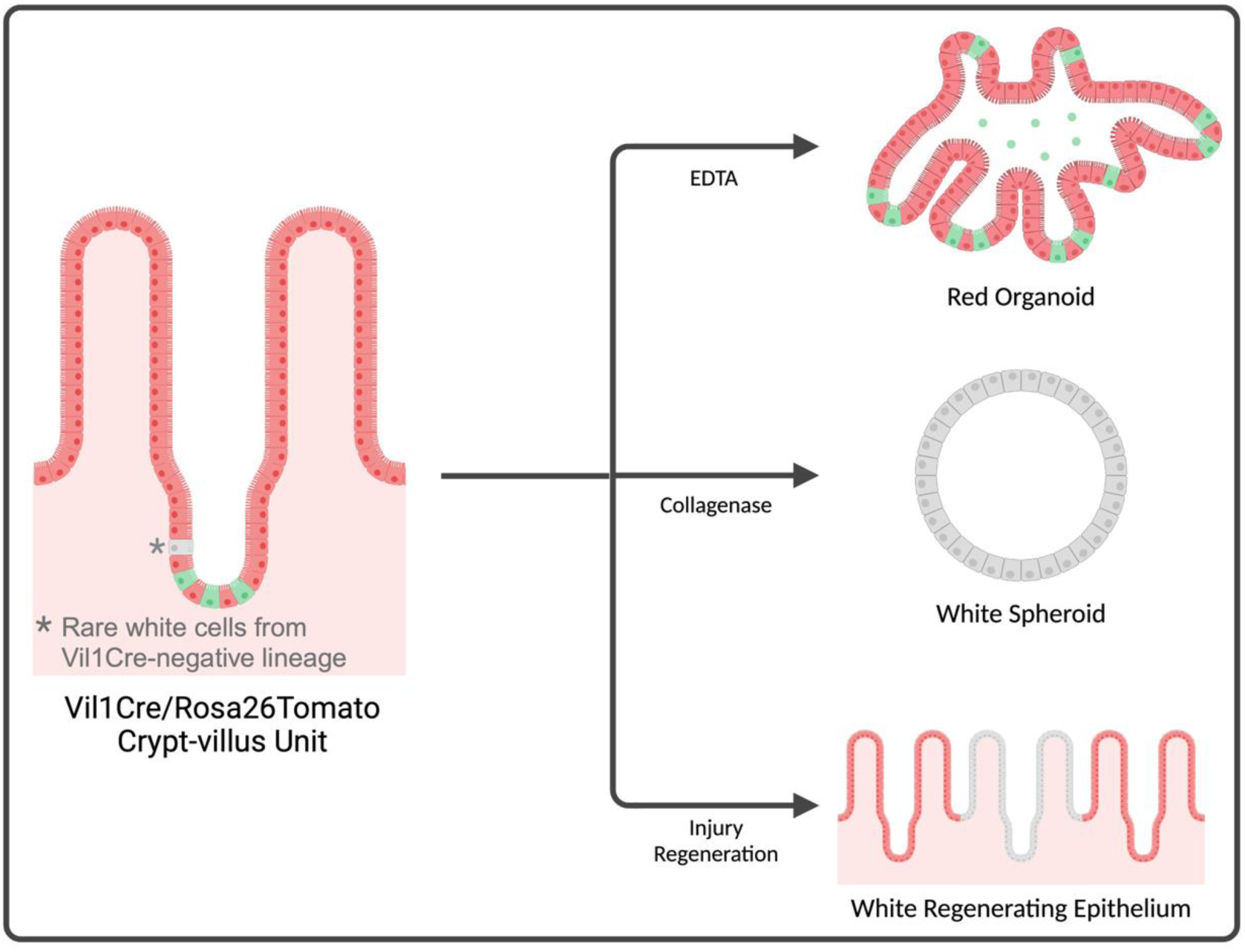

## Introduction

Maintenance of the intestinal epithelium involves actively cycling Lgr5-positive (Lgr5+ve) stem cells (<u>C</u>rypt <u>B</u>ase <u>C</u>olumnar cells, CBCs) located at the bottom of the crypts and taking care of cell renewing under homeostatic conditions (Barker et al., 2007; Beumer and Clevers, 2016). When submitted to harmful stresses causing cell death and/or affecting the integrity of the intestinal barrier, the epithelium triggers an extraordinary diversity of regenerative responses. These may involve activation of quiescent (“+4”) stem cells (Richmond et al., 2016; Yan et al., 2012), rerouting of early secretory progenitors to stemness (van Es et al., 2012), dedifferentiation of Paneth cells (Schmitt et al., 2018; Yu et al., 2018), enteroendocrine (Sei et al., 2018; Yan et al., 2017) or enterocyte progenitors (Tetteh et al., 2015) (for reviews, see (Bankaitis et al., 2018) (Capdevila et al., 2021)). When activated, these various mechanisms lead to reconstitution of the Lgr5+ve stem cell pool, ready to ensure post-injury maintenance of the epithelium (Metcalfe et al., 2014). On top of this extensive phenotypic plasticity, two-way interconversion of CBCs and +4 stem cells has been demonstrated, leading to the conclusion that a hierarchical stem cell model might not apply in the intestine (Clevers and Watt, 2018; Post and Clevers, 2019; Shivdasani et al., 2021; Takeda et al., 2011; Yousefi et al., 2017).

Recent studies have demonstrated that a characteristic common to most, if not all, regenerative responses in the gut epithelium is the (re)acquisition of a fetal-like intestinal phenotype (Ayyaz et al., 2019; Fernandez et al., 2016; Nusse et al., 2018; Yui et al., 2018), with expression of part of a genetic program first identified in fetal intestinal spheroids (Mustata et al., 2013). In all cases, the cells at the origin of the regenerating tissue were not formally identified but they were assumed to belong to the main Lgr5+ve lineage (Ayyaz *et al*., 2019; Yui *et al*., 2018).

In the present study, we show that stably replatable spheroids, resembling strikingly fetal spheroids, are generated from homeostatic adult intestine following dissociation of the tissue with collagenase/dispase. Lineage tracing experiments demonstrate that the cells at the origin of these spheroids belong to a separate developmental lineage, independent of Lgr5+ve CBCs, and are involved in regeneration of the epithelium following ablation of CBCs. Our results lead to the conclusion that a hierarchical stem cell model applies to regeneration of the intestinal epithelium, in addition to the plasticity model.

## Results

### Collagenase/dispase treatment of adult intestine releases spheroid-generating cells

Collagenase/dispase treatment of intestinal tissue is used to isolate mesenchymal cells and grow them in 2D cultures (Powell et al., 2011). When the same protocol was used, but with the resulting material seeded in matrigel under Sato culture conditions (Sato et al., 2009), we observed growth of a mix of minigut organoids and variable numbers of cystic spherical structures resembling fetal intestinal spheroids (Fordham et al., 2013; Mustata *et al*., 2013) **(Fig.1A)**. Since the standard EDTA-based protocol (Sato *et al*., 2009) only generates organoids, we reasoned that the spheroids must originate from collagenase/dispase digestion of the material normally discarded after EDTA treatment, from which most of the crypts have been removed **(Fig.S1A)**. Using a two-step protocol (see Methods and **Fig.1B**), we obtained bona fide minigut organoids from the EDTA fraction, as expected, and only spheroids, together with mesenchymal cells, from the collagenase/dispase fraction **(Fig.1B)**. Upon replating, these spheroids could be stably cultured free of mesenchymal cells **(Fig.1B)**. Morphologically similar spheroids are obtained when EDTA-derived organoids are co-cultured with mesenchymal cells or in the presence of medium conditioned by mesenchymal cells **(Fig.1C)** (Lahar et al., 2011; Roulis et al., 2020). Under these conditions, the switch of organoids towards spheroid morphology is in direct relation with the amount of mesenchymal cells present in the co-culture and factors released by them **(Fig.S1B)**. However, contrary to collagenase/dispase-derived spheroids, spheroids derived from organoids revert to *bona fide* organoids as soon as they are freed from mesenchymal cells, or upon withdrawal of the conditioned medium **(Fig.1C)**. Using the classical matrigel dome-embedding protocol (but see below), the yield of adult collagenase/dispase-derived spheroids was variable with, on average, one out of three mice being productive.

**Figure 1.**
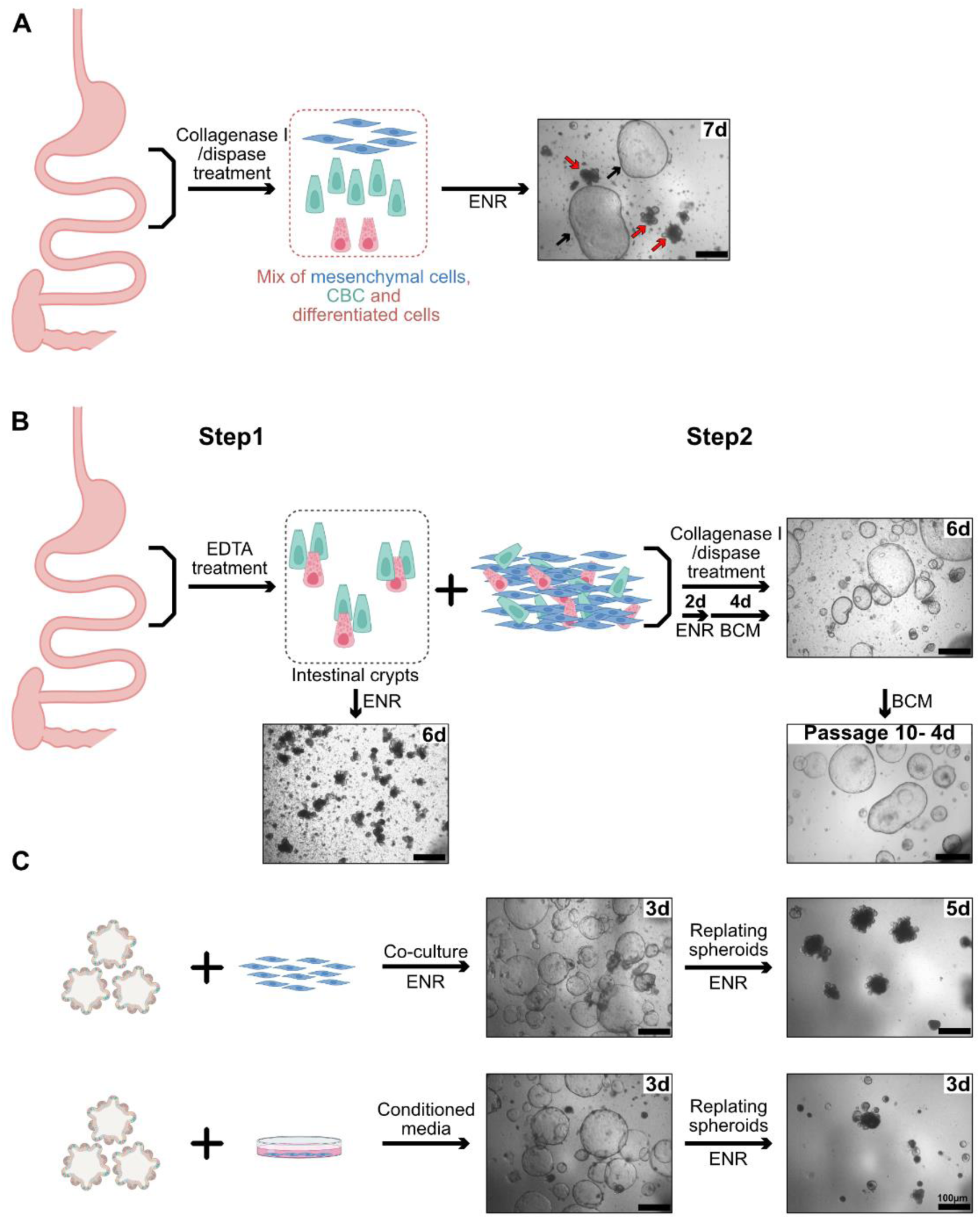
Collagenase/dispase treatment of adult intestine releases spheroid-generating cells. (A) Treatment of intestinal tissue with collagenase/dispase releases cellular material growing as spheroids (black arrows) when cultured in 3D, in addition to minigut organoids (red arrows). (B) EDTA treatment of intestinal tissue (step1) releases crypts growing only as organoids, whereas action of collagenase/dispase on the material remaining after EDTA treatment (step2) generates only spheroids that can be serially replated free of mesenchyme. (C) EDTA-derived organoids in co-culture with intestinal mesenchymal cells or in the presence of medium conditioned by intestinal mesenchyme convert to a spheroid phenotype but revert to the organoid phenotype in the absence of mesenchyme influence. All scale bars: 100 µm. Indications in the upper right corner of each picture refer to the days after initiation of cultures.

### Adult intestinal spheroids grow in absence of Rspondin/EGF/Noggin

The yield of fetal intestinal spheroids obtained by the EDTA Sato protocol decreases steadily during fetal life to reach zero shortly after birth (Fordham *et al*., 2013; Mustata *et al*., 2013). To explore the possibility that adult spheroids would be identical to fetal spheroids, but generated from cells more tightly attached to the matrix in postnatal tissue, we compared the culture medium requirement of both kind of spheroids. Whereas, in our hands, fetal spheroids grow best in ENR-containing medium (EGF, Noggin, Rspondin) (Sato *et al*., 2009), and depend totally for survival on Rspondin and expression of the Lgr4 gene (Mustata *et al*., 2013), adult spheroids survive and thrive in plain culture medium without addition of growth factors **(Fig.2A,B,C)**. This characteristic excludes that they could originate from pancreatic tissue possibly contaminating duodenal preparations (Huch et al., 2013). In long-term cultures, plain basal cell medium (BCM) allowed maintenance of the clear spheroid phenotype, whereas in EN (EGF, Noggin) or ENR they became dark and shrink **(Fig.2B,C)**, if not replated every two days. In addition, and contrary to fetal spheroids, Lgr4-deficient spheroids from P15 mice survive **(Fig.2D)** and could be replated, which is coherent with their independence from exogenous Rspondin. Similar intestinal spheroids displaying a Wnt (Rspondin)-independent phenotype have been obtained from culture of isolated Bmi+ve “+ 4” cells (Smith et al., 2018), or in models of intestinal regeneration, *in vivo* (Nusse *et al*., 2018) or *ex vivo* (Yui *et al*., 2018). These studies constituted the first indication of a potential link between adult collagenase/dispased-derived spheroids and tissue regeneration.

**Figure 2.**
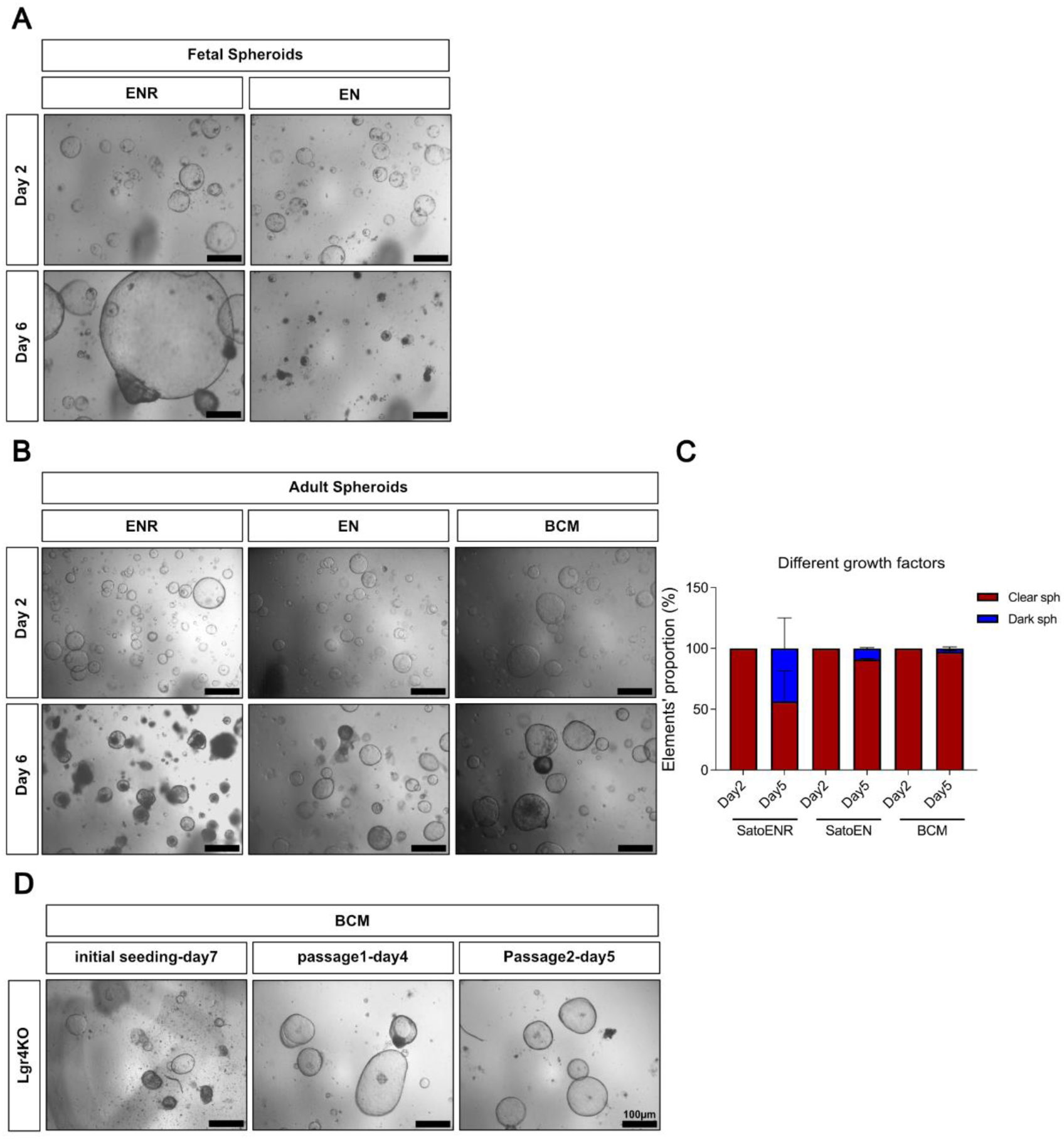
Adult intestinal spheroids grow in absence of EGF/Noggin/Rspondin (ENR). (A) Fetal spheroids derived from E15.5 embryonic intestine rely on Rspondin for their growth in vitro. (B) Adult spheroids can be cultured in the presence ENR medium as well as in EN medium but thrive better in plain medium devoid of growth factors (BCM). (C) the proportion of dark and clear spheroids at day 2 and day 6 after being in different culture media. (D) Lgr4KO spheroids derived from postnatal intestine (P15) grow normally and can be subcultured in BCM conditions. Scale bars: 100 µm.

### Adult spheroid are made of poorly differentiated intestinal epithelial cells with downregulation of CBC markers and expression of a regeneration program

Bulk RNAseq demonstrated that adult spheroids display partial downregulation of genes specific for most intestinal lineages (Haber et al., 2017) [enterocytes (Sis, Alpi, Treh…), Goblet cells (Clca3a1, Clca3a2, Clca3b, Muc2, Muc3, Fcgbp..), Paneth cells (Defa-rs1, Defa17, Lyz1, Gm15284, AY761184…), enteroendocrine cells (Chgb, Neurod1, Neurog3, Vwa5b2..)]**(Fig.3A, Fig.S2A-E, Table S1)**. However, they are clearly intestinal in origin with robust expression of several transcripts enriched in the intestinal epithelium (Vil1, Crip1, Ick, Epcam, Krt8, Krt19, Krt7) **(Fig.3B)**. Unexpectedly given their epithelial phenotype, with the notable exception of Vimentin, spheroids display strong upregulation of a set of genes expressed in intestinal mesenchyme (Li et al., 2007) **(Fig.3C)**, together with a matrisome gene signature (type 4 collagens, laminins, metalloproteases, fibronectin, Ctgf, cytokines, Pdgf….) (Naba et al., 2012) **(Fig.3D)**. All these characteristics were observed for spheroids cultured in ENR (that needed being replated every other day, see **Fig.2**), and more so in spheroids stably cultured in basal medium (**Table S2**).

**Figure 3.**
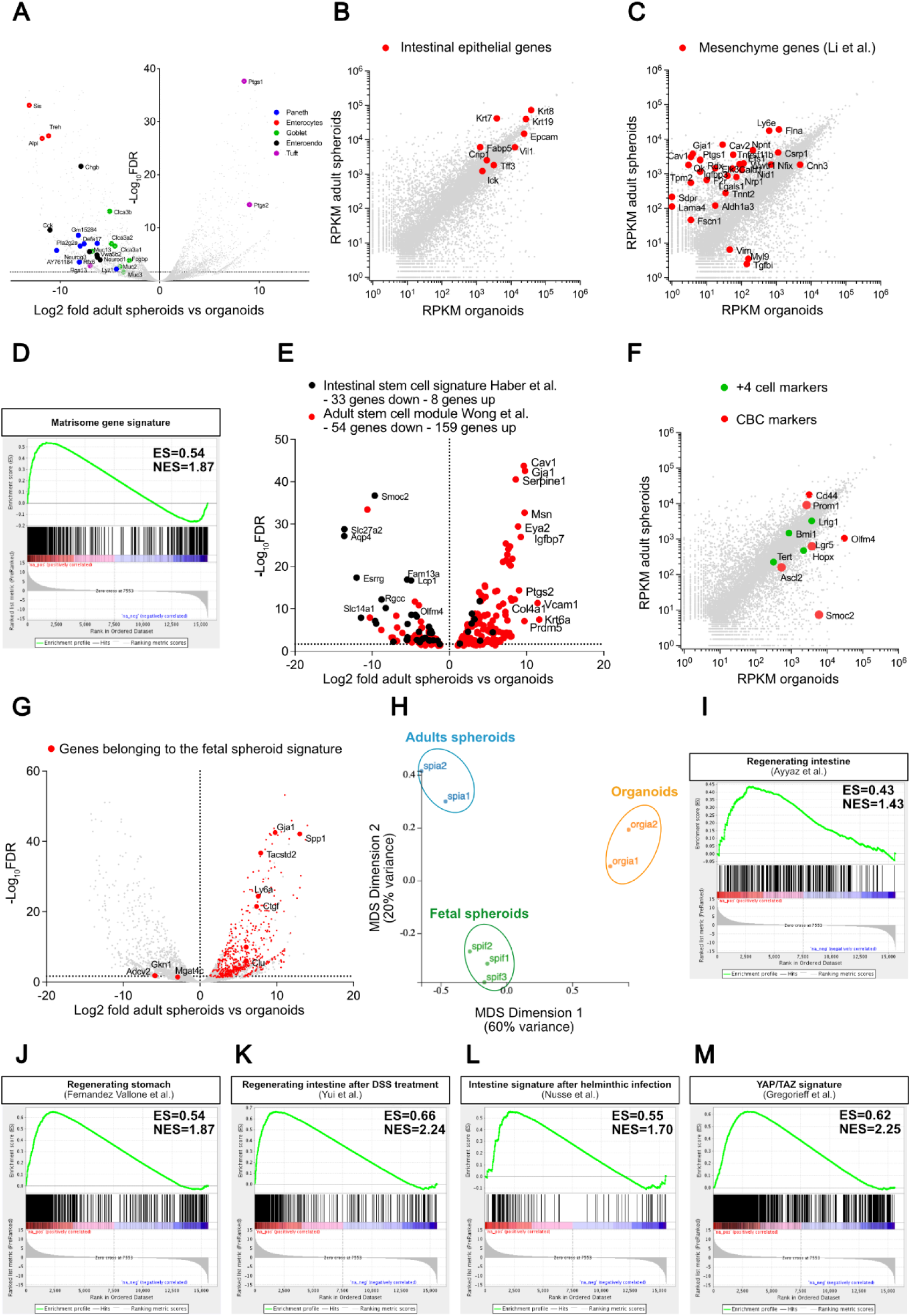
Gene expression program of adult spheroids. (A) Volcano plot showing that, compared to organoids, adult spheroid transcripts display partial downregulation of markers representative of all five intestinal differentiated cell types; to allow pertinent comparison spheroids and organoids were cultured in the same ENR-containing medium. (B) Logarithmic plot showing reads per kilo base per million (RPKM) measured for adult spheroids and organoids; red dots point to expression of genes illustrating the intestinal epithelial nature of spheroids. (C) Logarithmic plot showing RPKM for adult spheroids and organoids; red dots point to expression of genes belonging to the intestinal mesenchymal gene signature by Li et al (Li *et al*., 2007). (D) Pre-ranked gene set enrichment analysis (GSEA) showing positive correlation between adult spheroid up-regulated genes (versus organoids) and matrisome signature described by Naba et al. (Naba *et al*., 2012). (E) Volcano plot showing upregulation of an “adult tissue stem cell signature” (red dots= 54 genes down- and 159 genes up-regulated) (Wong et al., 2008) and down regulation of an “intestinal stem cell signature” (black= 33 genes down- and 8 genes up-regulated) (Haber *et al*., 2017) in adult spheroids, compared to organoids (total number of genes modulated with an FDR<0.05: 3329). (F) Logarithmic plot showing RPKM for adult spheroids and organoids; colored dots point to “+4” (green) or CBC (red) stem cell markers; with the exception of Prominin and CD44, CBC markers where downregulated and “+4 markers” were expressed at levels similar to those in organoids. (G) Volcano plot of adult spheroid versus organoid transcriptomes, showing upregulation of the vast majority of genes belonging to the fetal spheroid signature (red dots) (Fernandez *et al*., 2016). (H) Multidimensional scaling plot (MDS) displaying relatedness between adult intestinal spheroids (blue), fetal intestinal spheroids (green), and intestinal organoids (orange). (I-M) Pre-ranked gene set enrichment analyses (GSEA) showing positive correlation between adult spheroid up-regulated genes (versus organoids) and the gene signature of Clu+ve “revival stem cells” from irradiated intestine (Ayyaz *et al*., 2019) (I), regenerating stomach (Fernandez *et al*., 2016) (J), regenerating intestine induced by dextran sulfate sodium (DSS) treatment in a murine model of colitis (Yui *et al*., 2018) (K), the intestine following helminthic infection (Nusse *et al*., 2018) (L), and Yap/Taz gene expression signature (Gregorieff *et al*., 2015) (M). ES = enrichment score, NES = normalized enrichment score.

In agreement with their aptitude to be serially replated over long periods (26 replatings with maintenance of a stable phenotype) and to resist freeze/thaw cycles, adult spheroids express a set of upregulated genes overlapping significantly with an “adult tissue stem cell module” [159/721 genes; q value 2.11 e^-94^) **(Fig.S2F)**]. However, they paradoxically display downregulation of several genes defining the “intestinal stem cell signature”, including the paradigmatic CBC markers (Lgr5, Ascl2, Smoc2, Olfm4) **(Fig.3E-F; Fig.S2G)** (Haber *et al*., 2017). Markers of “+4” stem cells (Bmi1, Tert, Lrig1, Hopx) were all expressed in adult spheroids, with no significant up- or downregulation, compared to organoids **(Fig.3F)**(Richmond *et al*., 2016).

Bulk RNASeq data confirmed the close relationship of adult spheroids with fetal spheroids **(Fig.3G, Fig.S2H-J)**. Out of the 692 genes constituting a fetal intestinal and gastric spheroid signature (Fernandez *et al*., 2016), only three were slightly downregulated, and 533 were upregulated in adult spheroids, among which, Ly6a/Sca1, Tacstd2/Trop2, Gja1, Ctgf, Clu and Spp1 **(Fig.3G)**. However, despite their similarity, adult and fetal spheroids are clearly distinct **(Fig.3H)**, with adult spheroids showing 1,169 and 768 genes up- or downregulated, respectively, when compared to fetal intestinal spheroids **(Table S3)**. GSEA analyses for hallmarks comparing the adult and fetal spheroid transcriptomes, revealed highly significant upregulation of genes implicated in epithelial-mesenchymal transition (61 genes; FDR q value e^-51^), activation of the NFkB pathway (58 genes; FDR q value e^-48^) or inflammation (42 genes; FDR q value e^-28^) and stimulation by interferon gamma (39 genes; FDR q value e^-25^) **(Table S4)**.

Finally, GSEA analyses demonstrated striking similarities between the transcription program of adult spheroids and those observed in several models of regenerating gut (Ayyaz *et al*., 2019; Fernandez *et al*., 2016; Nusse *et al*., 2018; Yui *et al*., 2018) **(Fig.3I-L)**, with expression of a robust Yap/Taz gene signature (Gregorieff et al., 2015) **(Fig.3M)**.

### Adult spheroids originate from activation of a regeneration program in cells following their release from the extracellular matrix by collagenase/dispase

Since adult spheroids were obtained in variable amounts and not from all mice, we initially hypothesized that they originated from regions of the intestine of some animals in which regeneration takes place following minimal “physiological” injuries. However, using a new two-layer sandwich protocol described by the Lutolf group (Bues et al., 2020), we reproducibly obtained adult spheroids from all mice **(Fig.S3A-C)**. In this protocol, cells dissociated from the mesenchyme by collagenase/dispase are first allowed to spread on the surface of matrigel, before being covered by a second matrigel layer. Considering the known relation between matrix interaction and stem cell quiescence/activation (Gilbert et al., 2010; Machado et al., 2021; Montarras et al., 2005), our interpretation is that spreading of cells at the origin of spheroids on the first layer of matrigel might mimic more closely the effects of tissue injury than abrupt culture in matrigel. The spheroids obtained with the dome and sandwich protocols were very similar regarding transcriptome and culture medium requirements **(Fig.S3B-E)**.

In most regenerating conditions of the intestinal epithelium, after initial activation of a Yap/Taz genetic program, novel Lgr5-positive CBCs are generated. We propose that spheroids observed after collagenase/dispase treatment correspond to cells “frozen” in a stable Yap/Taz regenerating-like phenotype.

Together, these results suggest the existence of peculiar stem cells, tightly attached to extracellular matrix, which activate a regeneration program when they are severed from their normal environment by collagenase/dispase treatment and cultured in matrigel.

### Organoids and adult spheroids originate from separate developmental intestinal lineages

Given expression of both mesenchymal and epithelial markers in adult spheroids **(Fig.3B-D)**, we wanted first to ensure that they belong to the epithelial intestine lineage. To that aim, we used the Tg(Vil1Cre)997Gum/J mouse line (Madison et al., 2002), expressing a Cre recombinase transgene specifically in the intestinal epithelium, starting at E12.5. To our surprise, whereas close to 100% of EDTA-organoids and spheroids generated from E16.5 Vil1Cre/Rosa26 mice were positive for reporter expression, as expected **(Fig.4A)**, a vast majority of the adult spheroids were negative (70-90%) **(Fig.4B-D)**, despite displaying robust expression of Villin **(Fig.3B**, **Fig.4E)**. This difference is in strong contrast with the low level of mosaicism (at most a few percent) displayed by this mouse line (Madison et al., 2002), which we interpret as an indication that absence of Vil1Cre expression constitutes a potent tracing opportunity. To explore the causes behind this observation, we quantified Cre transcripts in spheroids and organoids from Vil1Cre/Rosa26Tomato mice. This demonstrated robust Cre expression in organoids and very low or absence of Cre mRNA in both “white” and “red” spheroids **(Fig.4E)**, the latter probably related to the exquisite sensitivity of the Rosa26 Ai9 Tomato reporter to traces of recombinase (Liu et al., 2013). Further confirming these results, direct measurement of recombination at the Rosa26Tom locus showed absence of recombination in white spheroids, when organoids were 100% recombined **(Fig.4F)**. In addition, no significant tracing of spheroids was obtained from Lgr5CreERT2/Rosa26Tom animals 5 days or 30 days after three pulses of Tamoxifen **(Fig.4G)**. This indicates that, while expressing low but detectable levels of Lgr5 upon culture **(Fig.3F)**, adult spheroids originate from a Lgr5-negative developmental linage.

**Figure 4.**
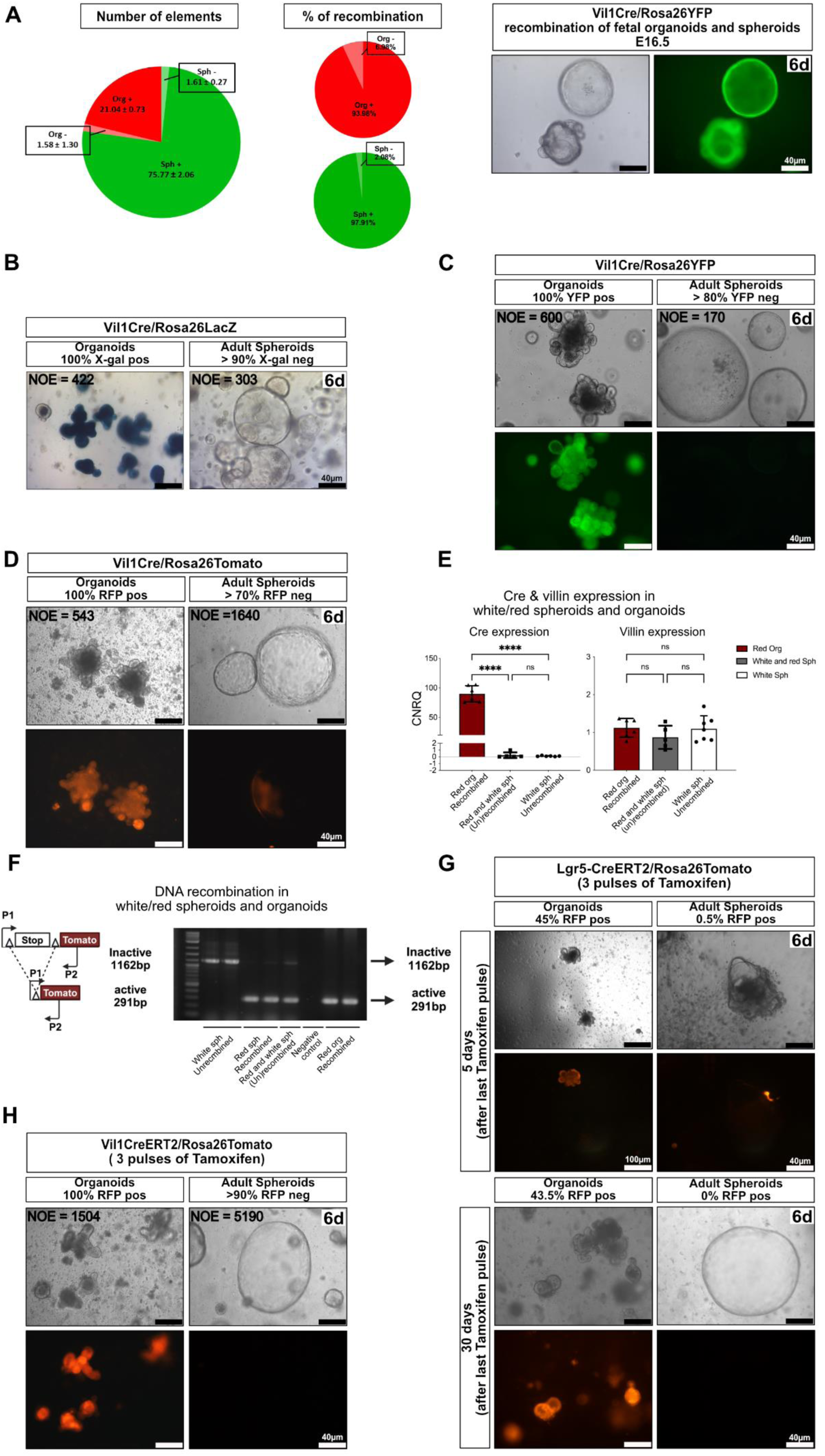
Organoids and adult spheroids originate from separate developmental intestinal lineages. (A) Proportion of elements and % of recombined fetal organoids and spheroids from E16.5 Vil1Cre/Rosa26YFP mice (left); representative pictures (right) at day 6 of initial seeding (n=3). (B-D) Representative pictures showing absence of recombination in spheroids from Vil1Cre mice using Rosa26LacZ (n=2) (B), Rosa26YFP (n=7) (C) or Rosa26Tom (n=4) (D) reporters 6 days after initial seeding. (E) qRT-PCR of Cre (expressed as CRNQ) and Villin (expressed relative to level in organoids) measured in organoids and in Tom+ve (red/recombined) and Tom-ve (white/unrecombined) spheroids. (F) PCR strategy targeting the Rosa26Tom locus showing absence of recombination in white spheroids prepared from Vil1Cre/Rosa26Tom mice. (G) Absence of recombination in spheroids derived from Lgr5CreERT/Rosa26Tom mice after three pulses of tamoxifen (n=2 for short chase n=3 for long chase). (H) Absence of recombination in spheroids cultured from Vil1CreERT2/Rosa26Tom mice after three pulses of tamoxifen (n=4). Scale bars: 40 µm, except in upper left pictures of panel F (100 µm). Indications in the upper right corner of each picture refer to the days after initiation of cultures. NOE= number of elements counted for each condition. Statistical analyses for panel E: one-way ANOVA test with Tukey’s multiple comparison tests; for Cre expression: ns p=0.9996, **** p<0.0001; for Villin expression: ns p > 0.05.

Since adult spheroids display robust expression of Villin **(Fig.3B)**, one must conclude that, contrary to the endogenous Villin promoter, the 12.4 Kb transgene fragment in the construct stays inactive in the whole cell lineage at the origin of adult spheroids. Similarly, tamoxifen administration to Vil1CreERT2/Rosa26YFP or Vil1CreERT2/Rosa26Tomato mice, which harbor a shorter Villin promoter fragment (9kb) (el Marjou et al., 2004), yielded also 100% recombined organoids and close to 100% un-recombined spheroids **(Fig.4H)**. These experiments suggest that transcription factors must be acting differently at the endogenous Villin gene promoter/enhancers in organoids and adult spheroids, respectively. Results from AtacSeq experiments give support to this hypothesis, showing different patterns of open chromatin in the vicinity of the endogenous Villin gene in spheroids and organoids **(Fig.S4)**. We interpret this as meaning that different ci-regulatory sequences are used for endogenous Villin gene expression in organoids and spheroids, respectively.

We conclude from the differential behavior of the Vil1Cre transgenes in adult spheroids and organoids that the two structures originate from different developmental lineages.

### Adult spheroid-generating cells are involved in regeneration of the intestinal epithelium following injury

Since non-recombined cells are exceedingly rare in the intestinal epithelium of Vil1Cre/Rosa26Tom mice (Madison *et al*., 2002) under homeostatic conditions, our results suggest that collagenase/dispase treatment of intestinal tissue followed by 3D culture in matrigel triggers proliferation of rare quiescent cells belonging to a hitherto undescribed intestinal lineage. Collagenase/dispase solution would act as a proxy for injury and adult spheroids would correspond to a stable avatar of intestinal regenerating cells, trapped in a state compatible with unlimited ex-vivo culture. If this hypothesis holds true, we reasoned that we could use absence of recombination in Vil1Cre/Rosa26Tom intestine as a tool to trace the possible involvement of adult spheroid-generating cells in intestinal regeneration. We tested this hypothesis in two injury models.

In the first, the ubiquitously expressed anti-apoptotic gene Mcl1 was ablated in the intestinal epithelium of Vil1CreERT2/Mcl1^fl/fl^ mice (Vikstrom et al., 2010a) **(Fig.5A)**. A wave of apoptosis was observed in the crypts of animals treated with tamoxifen, with a maximum peaking at 24h **(Fig.5B,C)**. When the tissue was processed for organoid and spheroid production, no organoids could be obtained from Tamoxifen-treated animals at 24h, indicating effective ablation of CBCs. Unexpectedly, rare spheroids were cultured from the EDTA fraction at 24 h after Tamoxifen. Spheroids were obtained from collagenase/dispase fractions irrespective of Tamoxifen treatment **(Fig.5D,E)**, which fits with the hypothesis that they originate from cells belonging to a lineage different from CBCs. Looking for regenerating crypts possibly growing from this non-recombined lineage, we performed RNAscope in situ hybridization (ISH) with two Mcl1 probes, which yielded unexpected results. Removal of the floxed segment of the Mcl1 gene resulted in a strong increase of the ISH signals obtained in the intestinal epithelium with both probes **(Fig.S5)**. This must result from the removal of a transcriptional silencer from the gene, or from stabilization of the mRNA transcribed from the recombined gene (or both). Whatever the explanation, this increase in ISH signal constitutes a sensitive and convenient proxy for recombination, allowing to test for the presence of un-recombined regenerating crypts. When performed 16h, 24h, 36h and 60h after tamoxifen administration to Vil1CreERT2/Mcl1^fl/fl^ mice, Mcl1 ISH showed a wave of un-recombined cells progressing from the crypts towards the villi **(Fig.5F,G)**. Given the absence of organoid-generating cells in tamoxifen treated Vil1CreERT2/Mcl1^fl/fl^ mice at 24h **(Fig.5D,E)**, it seems that recombination efficiently ablates most CBCs and that, indeed, regeneration starts from un-recombined cells. However, in this model, we cannot differentiate between the hypotheses that the wave of un-recombined crypts would originate from CBC or TA cell escapers rather than from a specific lineage. We therefore turned to a second model of injury involving constitutive Vil1Cre expression.

**Figure 5.**
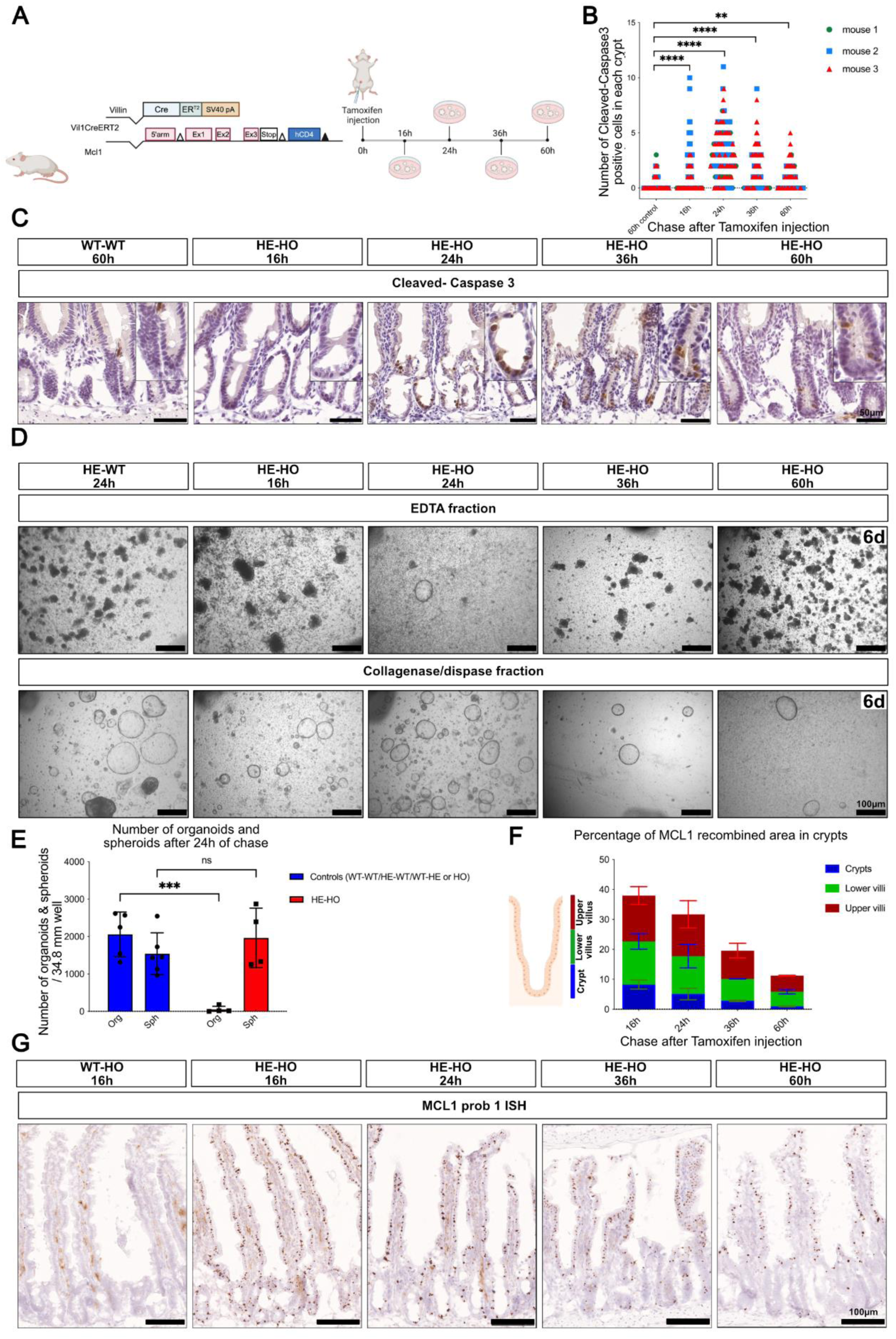
Regeneration of Mcl1-ablated intestine originates from cells escaping recombination triggered by VilCreERT2. (A) Schematic summary of the experiment. (B) Number of caspase 3-positive cells in crypt cells at various time after tamoxifen injection (n=3). (C) Representative pictures of crypts with Caspase 3-positive cells (WT/WT: wild type mice; HE/HO: Vil1CreERT2 heterozygotes, homozygote for Mcl1^fl/fl^). (D) Representative pictures showing the yield of organoids (EDTA fraction panels) or spheroids (Collagenase-dispase fraction panels) at various time points after tamoxifen administration. Cultures are shown 6 days after plating. (E) Quantification of spheroids and organoids per well, 24 hours after Tamoxifen injection; each symbol illustrates results from one mouse (n>4). (F) Percentage of epithelial surface containing Mcl1^fl/fl^ recombined cells, along the crypt-villus axis at various times after tamoxifen administration, n=2 to 7 mice (see also Fig.S5). (G) Representative pictures of intestinal epithelium showing Mcl1 RNAscope signals at various times after tamoxifen administration. Paradoxically, recombined cells show strong Mcl1-RNAscope signals (see text and Fig.S5). Statistical analyses: (B) Two-way ANOVA test with Dunnett’s multiple comparisons test: ** p= 0.0028; **** p<0.000. (E) Two-way ANOVA test with Šídák’s multiple comparisons: ns p>0.05; *** p<0.001. Scale bars: 40 μm (C) and 100 μm for (D,G).

In this model, CBCs are ablated by diphtheria toxin (DT) administration to mice expressing the human diphtheria toxin receptor gene under control of Lgr5 regulatory regions (Lgr5-DTR knock-in mice (Tian et al., 2011)) **(Fig.6A)**. After three pulses of toxin administration over a period of 5 days, Vil1Cre/Lgr5-DTR/Rosa26Tomato triple heterozygotes mice lost weight **(Fig.6B)** and multiple Tomato-negative clones were observed all along the small intestine, some of which extending to the tip of villi and encompassing morphologically identified Paneth and goblet cells **(Fig.6C-F)**. Despite being Tomato-negative (i.e. Vil11Cre-negative), and similar to adult spheroids, these regenerating clones expressed Villin strongly in villi. At the crypt level, Villin expression was low irrespective of Tomato expression **(Fig.6G)**. Tomato-negative crypts were virtually absent in Vil1Cre/Rosa26Tom double heterozygote mice treated with the toxin **(Fig.6D,E)**. Unexpectedly, untreated Vil1Cre/Lgr5-DTR/Rosa26Tomato triple heterozygotes displayed more un-recombined crypts than treated Vil1Cre/Rosa26Tom double heterozygotes, although on average three times less than treated ones **(Fig.6D)**. This suggests that either monoallelic expression of Lgr5, or ectopic expression of the diphtheria toxin receptor in CBCs causes a low-grade chronic injury. The absence of increase of Tomato-negative crypts in two lines of mice with monoallelic expression of Lgr5 in CBCs (**Fig.S6A-D)** points to a direct effect of ectopic DTR expression. Although neglected by the many groups using DTR-expressing mice, an effect of HB-EGF (the DTR) has been documented in the intestine where it causes inflammation and stimulates cell proliferation (Chen et al., 2010). In line with this hypothesis, bulk RNAseq demonstrates activation in untreated Lgr5-DTR mice of a program overlapping that of adult spheroids **(Fig.S6)**, with a proportion of genes commonly up- or downregulated, belonging to Wong adult tissue stem cell module ((Wong et al. 2008) and **(Fig.3E)**) or differentiated intestinal epithelium, respectively **(FigS6E-I)**.

**Figure 6.**
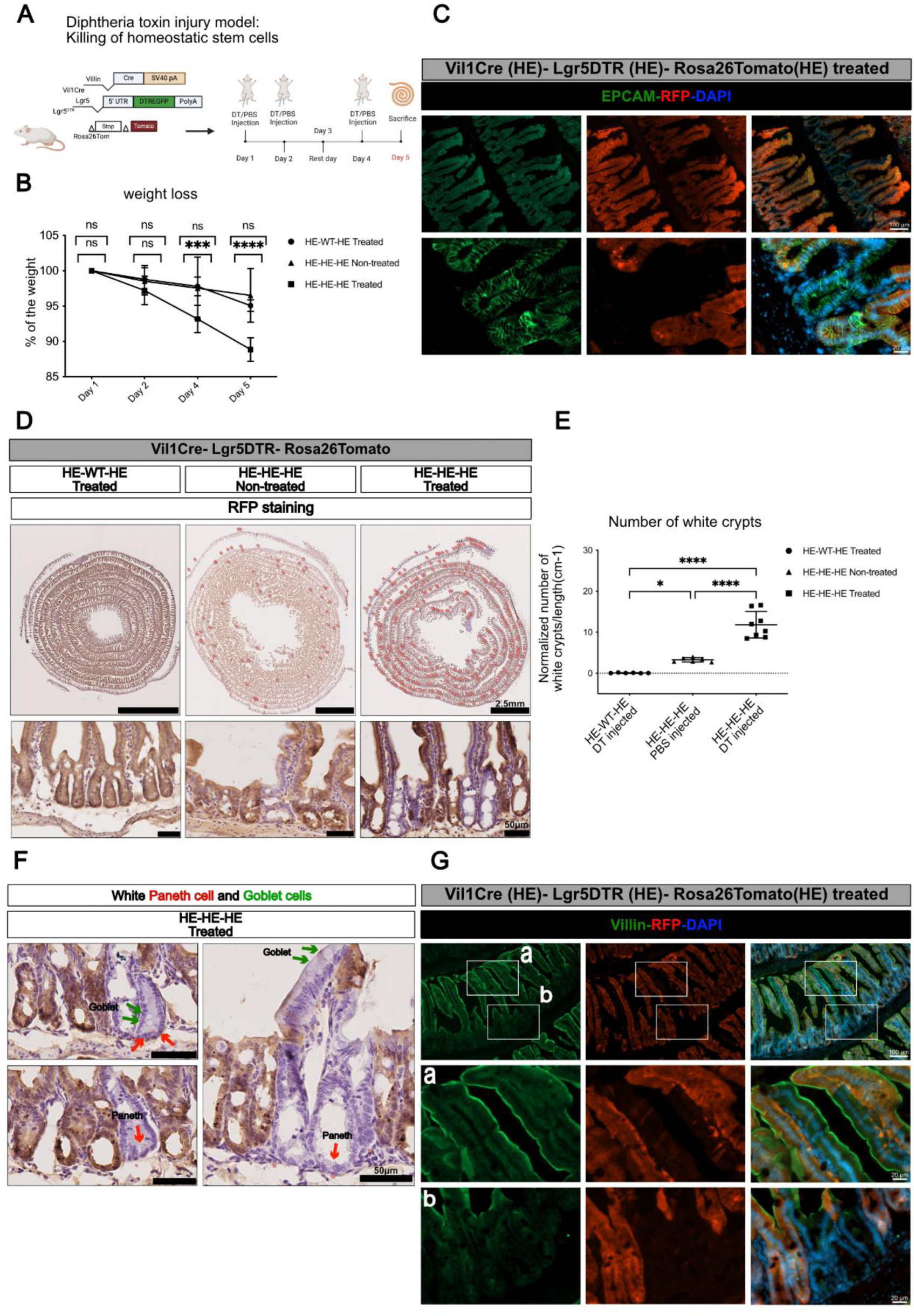
Contribution of VilCre-negative cells to epithelial regeneration following CBC ablation. (A) Schematic summary of the experiment. (B) Kinetics of weight loss of animals after DT treatment (n=6 to 8 mice in each group). (C) Immunofluorescence showing an RFP-negative (un-recombined) patch of epithelium in the small intestine of a Vil1Cre/Lgr5DTR/Rosa26Tom mouse treated with diphtheria toxin. (D-E) Quantification of RFP negative crypts in Swiss-rolls from animals treated or not with diphtheria toxin; HE-WT-HE: VilCre/Rosa26Tom double heterozygotes; HE-HE-HE: Vil1Cre/Lgr5DTR/Rosa26Tom triple heterozygotes. The crypts appearing as “white” after RFP immunohistochemistry (lower row) are marked on the rolls (upper row), counted and the results normalized to the length of the roll. Each symbol illustrates results from one animal (E); all samples were from mice sacrificed at day 5 (see panel A). (F) Higher magnification showing presence of “white” Paneth (red arrows) and Goblet cells (green arrows) in un-recombined epithelium patches. (G) Immunofluorescence showing Villin expression in villi (a) or crypts (b) of an RFP-negative (un-recombined) patch of epithelium in the small intestine of a Vil1Cre/Lgr5DTR/Rosa26Tom mouse treated with diphtheria toxin. Statistical analyses: (B) Two-way ANOVA test with Dunnett’s multiple comparisons test: ns p>0.05; *** p<0.001; **** p<0.0001. (E) one-way ANOVA test with Tukey’s multiple comparison tests: ns p>0.05; * p=0,0430; *** p<0.001; **** p<0.0001. Scale bars: 50 µm (C, D-bottom row, F, G) and 2.5 mm (D, upper row).

The kinetics of appearance of newly formed un-recombined (“white”) crypts was studied after a single pulse of DT **(Fig.7A)**. This demonstrated an increase at 48 hours, with further increase at day 10 and stable maintenance at day 30. The presence of newly formed white crypts one month after toxin administration indicates that the Vil1Cre-negative lineage is developmentally stable and does not turn on the transgene during differentiation of the various epithelial lineages occurring after regeneration (**Fig.7B)**. The relationship between white crypt generation and appearance of Clu-positive revival cells (Ayyaz *et al*., 2019) was then explored. In agreement with others and similar to what happens in the irradiation model, (Ayyaz *et al*., 2019; Yuan et al., 2023) Clu-positive cells were rare in crypts of untreated mice and their number transiently increased forty-eight hours after a single pulse of DT, and more so after three pulses of DT **(Fig.7C,D)**. Clu-positive cells were less frequently observed in white crypts (see “Total” versus “White” in **Fig.7C)**. This fits with the hypothesis that Clu expression marks acutely regenerating crypts and that a proportion of the white crypts not showing Clu expression corresponds to those that underwent regeneration at an undetermined time in the past, due to DTR expression in CBCs. This observation indirectly suggests that LGR5-DTR is not expressed in these “chronically white” crypts.

**Figure 7.**
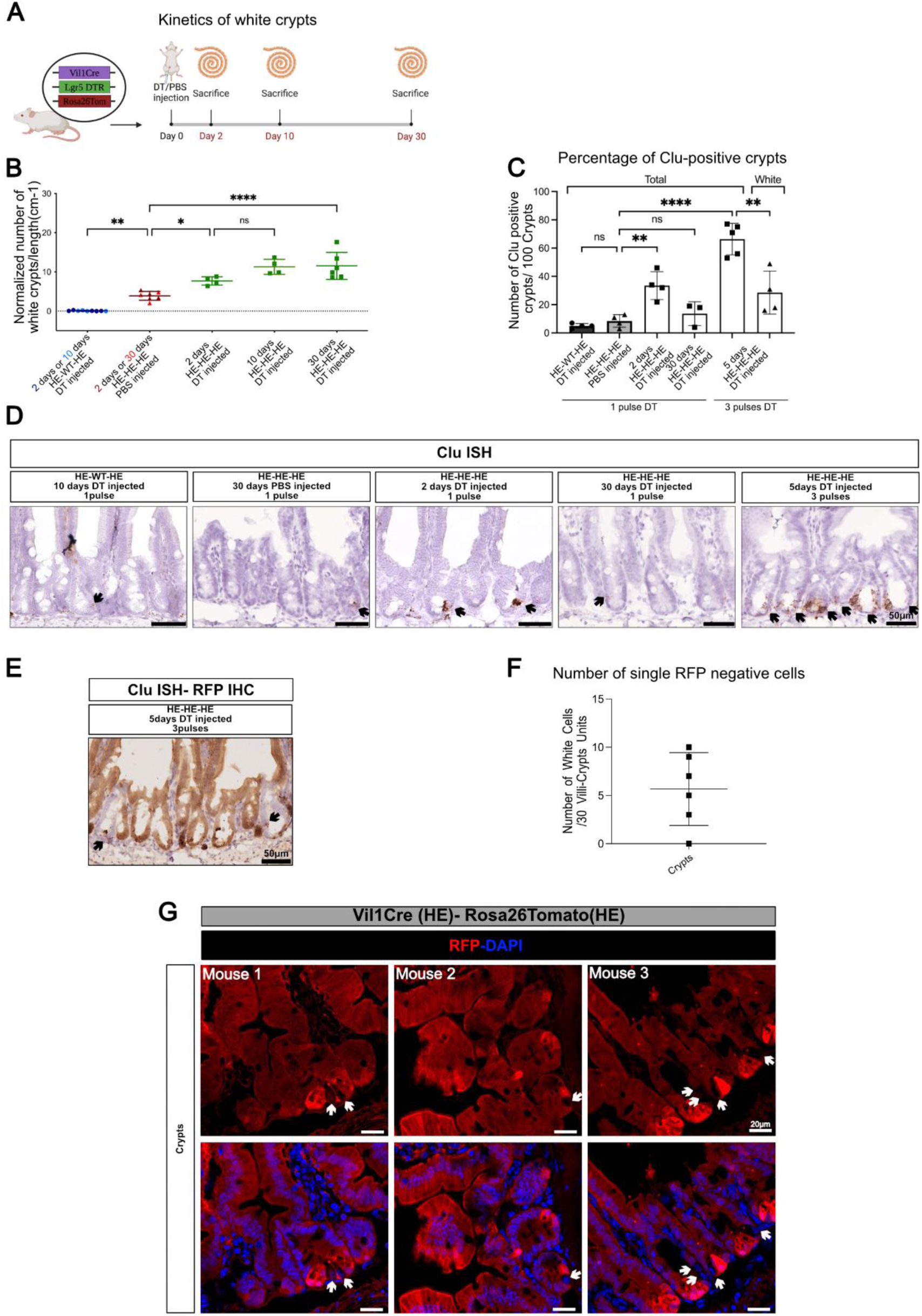
Kinetics of un-recombined crypt generation following CBC ablation. (A) Schematic summary of the experiment. Kinetics of unrecombined (white) crypts generation after a single injection of diphtheria toxin; HE-HE-HE, Vil1Cre/Lgr5DTR/Rosa26Tom triple heterozygotes; HE-WT-HE, Vil1Cre/Rosa26Tom double heterozygotes. Each symbol illustrates results from one animal. (C) Percentage of total or unrecombined (white) Clu-positive crypts by RNAscope ISH at different time points after 1 pulse (see panels A,B) or 3 pulses of Tamoxifen (see figure 6A,E). (D) Representative pictures of intestinal epithelium showing presence of Clu-positive cells in crypts after Tamoxifen administration at different time points. (E) Representative pictures of intestinal epithelium showing presence of Clu-positive cells in red and white crypts. (F) Number of single Tomato-negative cells in 30 crypts in Vil1Cre/Rosa26Tom double heterozygote mice; each symbol illustrates results from one mouse (n=6). (G) Representative pictures of immunofluorescence of Tomato-negative single cells in crypts. Statistical analyses (panels B,C): one-way ANOVA test with Tukey’s multiple comparison tests; ns p>0.05; *p<0.02; **p<0.01; **** p<0.0001.

Altogether, our results strongly suggest that the same Vil1Cre-negative cell lineage is at the origin of adult spheroids and a fraction of regenerating crypts observed in the diphtheria toxin model of injury. The actual proportion of crypts regenerating from this lineage is difficult to evaluate. Indeed, given the extreme sensitivity of the Rosa26Tom reporter to traces of recombinase [(Liu *et al*., 2013) and **Fig.4**], it is likely that using absence of recombination as a criterion leads to an underestimation of their number.

### Cells at the origin of the Vil1Cre-negative lineage are quiescent Olfm4-positive cells

Un-recombined cells are exceedingly rare in crypts of Vil1Cre/Rosa26Tomato double heterozygotes **(Fig.7F,G)**. In an attempt to characterize them, Epcam+ve/Rosa26Tomato-negative cells (“PlusMinus” cells, hereunder) were purified by FACS from the tissue fraction giving rise to adult spheroids **(Fig.8A**). This fraction is made of what remains after EDTA extraction of crypts from the total intestine (see **Fig.1B**). Microscopic inspection reveals that it is essentially made of mesenchyme delineating ghosts of crypts **(Fig.S1A)**. Effectively, PlusMinus cells represented a small proportion of the cells in this fraction (0.6 %, **Fig.8A**), which were then subjected to scRNAseq, together with Epcam-positive/Rosa26Tom-positive cells from the same fraction (“PlusPlus” cells, hereunder).

**Figure 8.**
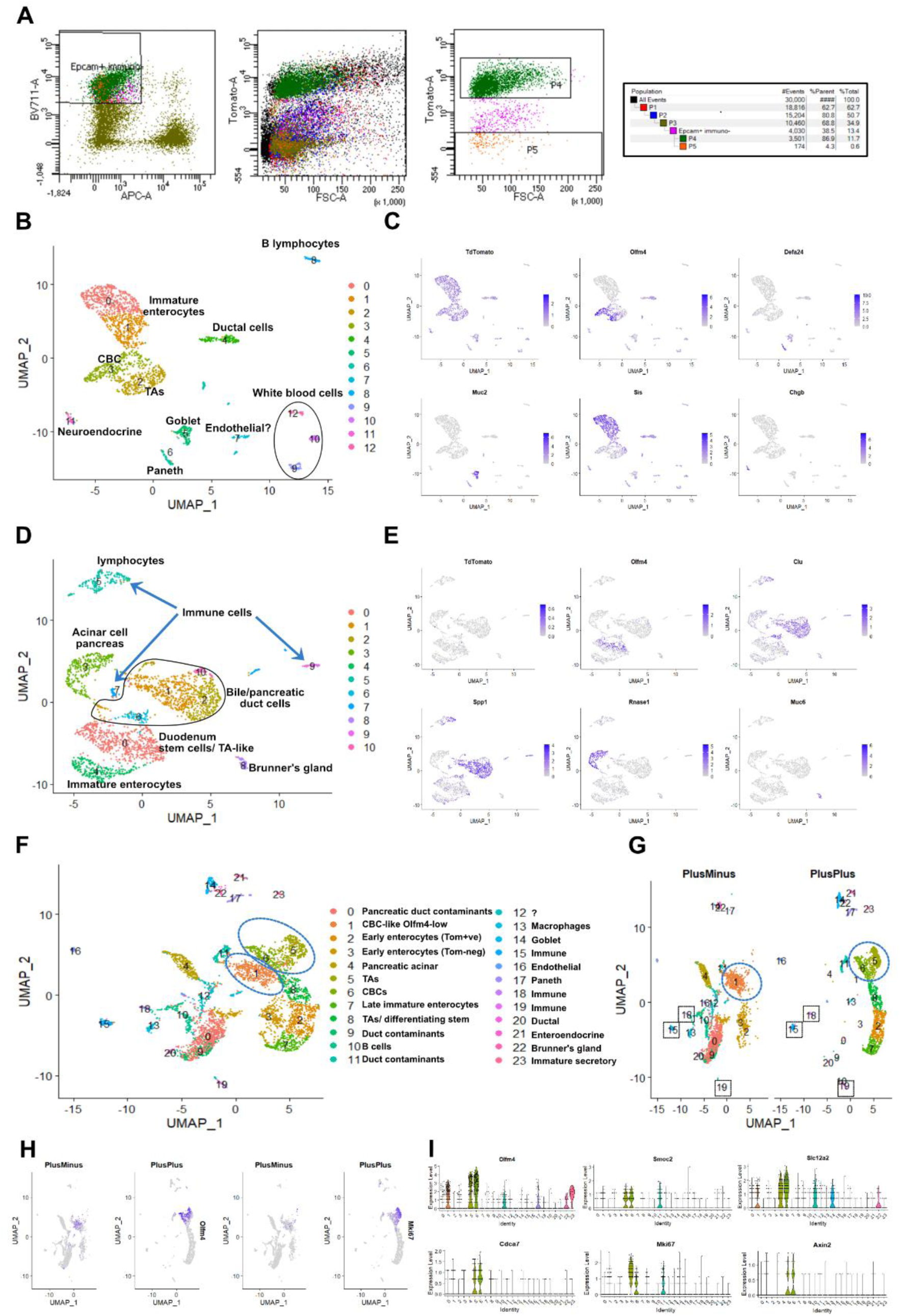
scRNAseq of Epcam-positive-Rosa26Tom-positive (PlusPlus) and Epcam-positive-Rosa26Tom-negative cells (PlusMinus) (A) Fluorescent sorting of PlusPlus cells (P4 window) and PlusMinus cells (P5 window). (B) UMAP representation of scRNAseq of 2154 PlusPlus cells showing 12 clusters identified by their differentially expressed genes (DEGs). (C) Same UMAP showing expression of Tdtomato, Olfm4, Defa24, Muc2, Sis and Chgb. (D) UMAP representation of scRNAseq of 2650 PlusMinus cells showing 10 clusters identified by their DEGs. (E) Same UMAP showing expression of Tdtomato, Olfm4, Clu, Spp1, Rnase1 and Muc6. (F) UMAP representations of similar number of cells from PlusPlus and PlusMinus samples after merging data analysis in Seurat, with indication of the cell types identified from their DEG lists; Olfm4-positive clusters are indicated by dotted circles. (G) Same data displayed according to their PlusPlus or PlusMinus origin, with indication of overlapping contaminants (squares). (H) Same UMAP showing expression of Olfm4 and Mki67. (I) Violin plots showing expression level of canonical transcript of CBCs (Olfm4, Smoc2, Slc12a2, Cdca7), the Wnt target gene Axin2 and Mki67 in all the clusters involved in merged data analysis.

PlusPlus cells originate likely from crypts that were not released from their mesenchymal niche by EDTA. The Uniform Manifold Approximation and Projection (UMAP) plot of 2,154 PlusPlus cells displayed the expected TdTomato-positive clusters of CBCs (cluster 2, in **Fig.8B-C**), TAs, Paneth cells, precursors of enterocytes, Goblet and enteroendocrine cells in addition to minor contaminants (**Fig.8B,C**). The UMAP plot of 2,650 PlusMinus cells was more complicated to interpret due to the presence of more contaminants (**Fig.8D**) : (i) Epcam-positive TdTomato-negative cells originating from Brunners glands (Muc6-positive cluster 8) and (ii) pancreatic tissue sticking to the duodenum (Clu-positive, Spp1-positive ductal cells, clusters 1,2,6,10; **Fig.8D,E**); (iii) Epcam-negative cells having escaped FACS selection, coming from pancreatic acini (Rnase1-positive cluster 3) and (iv) immune cells (clusters 5,7 and 9) (**Fig.8D,E**). Nevertheless, the major cluster of PlusMinus cells (cluster 0) presented characteristics of epithelial duodenal stem cells, being Olfm4-positive and displaying abundant ribosomal protein transcripts (**Fig.8D,E** and **Fig.9A,B**). In order to avoid normalization biases and compare more accurately transcriptomes of PlusPlus and PlusMinus cells, we merged data from the two samples and analyzed the resulting UMAP (**Fig.8F,G**). The presence of common contaminants (e.g. clusters 15,18 and 19, **Fig.8G**) demonstrates absence of significant batch effects in the merged analysis. If we concentrate on Olfm4-positive clusters (stem cell-like, see dotted circles), we observe that those originating from PlusMinus (cluster 1 in the merged plot) and PlusPlus samples (clusters 6 (CBCs) and 5 (TAs)) do not overlap **(Fig.8F,G**). Several canonical transcripts of CBCs, including Olfm4 (Smoc2, Slc12a2, Cdca7), are present but strongly downregulated in cluster 1, together with the Wnt target gene Axin2 and Mki67 (**Fig.8H,I**). This suggested that cells in cluster 1 are related, but different from CBCs, and might be quiescent. Direct comparison of genes differentially expressed between cluster 1 and clusters 4 and 6 confirmed this hypothesis with downregulation of genes controlling cell cycle and mitosis (Mcm3,5,6,7, Cdk4, Ccnd2, Stmn1, ….) (**Fig.9C,D**).

**Figure 9.**
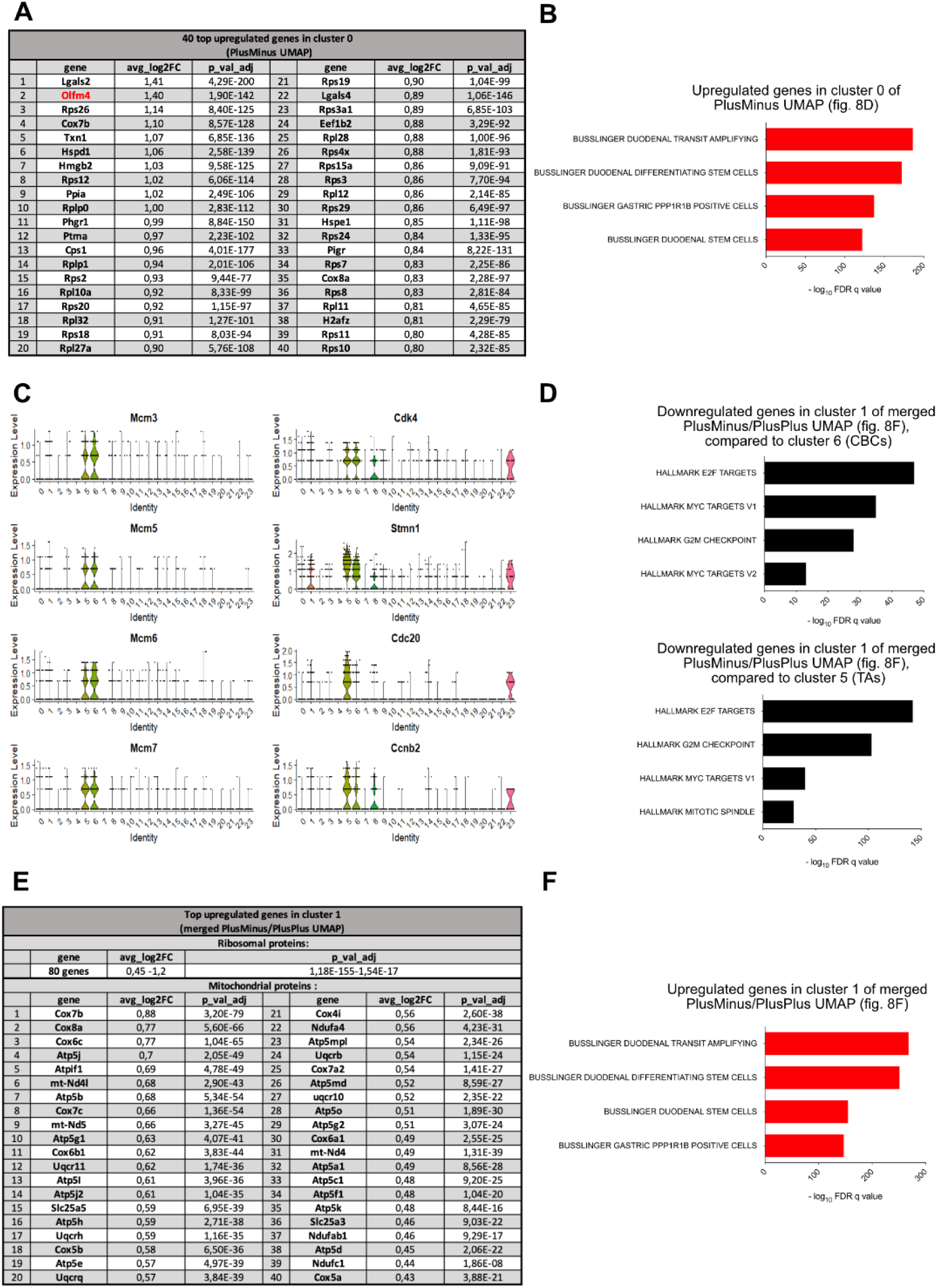
Cells at the origin of the VilCre-negative lineage are quiescent Olfm4-positive cells. (A) Table showing 40 upregulated genes in cluster 0 of PlusMinus (Fig.8D) sorted by log2 fold change. (B) GSEA MolSig results for biological cell types (C8) associated with the upregulated genes in cluster 0. (C) Expression levels of various genes implicated in cell cycle and mitosis across merged datasets (Fig.8F). (D) GSEA MolSig (hallmarks) results related to downregulated genes in cluster 1 of the combined PlusPlus and PlusMinus merged datasets (Fig.8F), in comparison to cluster 6 (CBCs) and cluster 5 (TAs). (E) Table showing 40 upregulated genes in cluster 1 of merged datasets (Fig.8F). (F) GSEA MolSig results for biological cell types (C8) associated with the upregulated genes in cluster 1 of the merged PlusPlus and PlusMinus datasets (Fig.8F).

From these results, we conclude that the Vil1Cre-negative lineage at the origin of adult spheroids and cells contributing to epithelial regeneration is made of quiescent stem cells sharing some characteristics with CBCs. The search for specific markers that would allow identification of these cells in the homeostatic intestinal epithelium yielded no results. Inspection of the differentially upregulated genes in these cells [cluster 0 of the PlusMinus UMAP (**Fig.9A**), or cluster 1 of the merged UMAP (**Fig.9E**)] revealed mainly the list of ribosomal protein genes characteristic of CBCs and, unexpectedly for quiescent cells, genes encoding cytochrome oxidase subunits, mitochondrial ATP synthase subunits and ATPase inhibiting factor (**Fig9A,E**).

## Discussion

In vertebrates, steady state maintenance and regeneration of tissues after injury is believed to follow two possible models: (i) a hierarchical model involving rarely dividing stem cells that generate progenitors with specific differentiation potentials (e.g. in hematopoiesis)(Seita and Weissman, 2010); (ii) a plasticity model characterized by the ability of committed or differentiated cells to de-differentiate, proliferate and reconstitute the various cell types (Wells and Watt, 2018). In the mammalian intestine, after initial vogue of a hierarchical model involving +4, rarely dividing label-retaining cells (Marshman et al., 2002), the identification of crypt base columnar cells (Cheng and Leblond, 1974) as actively dividing Lgr5+ve stem cells (Barker *et al*., 2007) changed the view. While Lgr5+ve cells were shown to be dispensable for daily epithelium maintenance (Tian *et al*., 2011), they are obligate actors of epithelial regeneration following e.g. irradiation (Metcalfe *et al*., 2014). According to current views, when the intestine is exposed to harmful stress, +4 quiescent cells, but also transit-amplifying progenitors or differentiated cells acquire, or re-acquire, Lgr5+ve stem cell characteristics and reconstitute the epithelium. In addition, reciprocal interconversion of +4 and Lgr5+ve cells has been demonstrated, leading to a model characterized by extensive two-way differentiation plasticity (Clevers and Watt, 2018; Yousefi *et al*., 2017).

There is no question that all the above phenomena can take place following epithelial damage, as they were convincingly demonstrated by lineage tracing experiments. However, their co-existence in-, or relative contribution to a given regenerating situation have not been assessed (Bankaitis *et al*., 2018; Buczacki, 2019). A role of Bmi1+ve cells in regeneration post-irradiation has been firmly established (Yan *et al*., 2012), but their heterogeneity and relation with Hopx+ve and Lgr5+ve cells (Beumer and Clevers, 2016; Li et al., 2016; Yousefi *et al*., 2017) made it difficult to identify precisely the cell(s) at the origin of the regeneration process. Our results identifying a cell lineage at the origin of stable adult spheroids produced after collagenase/dispase dissociation of intestinal tissue casts new light on intestinal regeneration. These cells are not traced by a Vil1Cre transgene that is expressed from E12 in the main Lgr5-positve epithelial lineage, nor by Lgr5CreERT. Spheroids similar to those described in our study have been obtained from the intestine of mice infested by helminths (Nusse *et al*., 2018), or in an *ex vivo* model of tissue regeneration (Yui *et al*., 2018). This strongly suggests that they represent avatars of regenerating cells, “frozen” in a stable, culture-compatible state. This conclusion is strengthened by the expression of a common fetal-like genetic program (Mustata *et al*., 2013) and a strong Yap/Taz signature, both characteristics reportedly associated with intestinal regeneration (Moya and Halder, 2019; Yui *et al*., 2018), and by the similarity of their transcriptome with that of “revival stem cells” (Ayyaz *et al*., 2019). Following ablation of CBCs in the diphtheria toxin injury model, Vil1Cre-negative crypt-villus units transiently expressing the revival cell marker Clu (Ayyaz *et al*., 2019) are generated, demonstrating the implication of this novel intestinal lineage in intestinal regeneration.

Constitutive ablation of the anti-apoptotic gene Mcl1 in the intestinal tract of Vil1Cre/Mcl1^fl/fl^ mice has been shown to cause chronic apoptosis and inflammation in crypts, with associated hyperproliferation and dedifferentiation leading to development of malignant tumors (Healy et al., 2020). A similar, but acute, inflammatory phenotype was observed in tamoxifen treated Vil1CreERT2/Mcl1^fl/fl^ mice. Interestingly, whereas the phenotype was homogenous in the induced mouse model (Vil1CreERT2), it was strongly mosaic in the constitutive version (Vil1Cre), with areas of normal histology keeping expression of the Mcl1 gene (Healy *et al*., 2020). Since both the Vil1Cre and Vil1CreERT2 transgenes are reportedly expressed throughout the intestinal epithelium (el Marjou *et al*., 2004; Madison *et al*., 2002), this observation is compatible with our hypothesis that a Vil1Cre- (or Vil1CreERT2)-negative intestinal lineage could be at the origin of regenerating intestinal cells. Our observation of a wave of un-recombined cells following tamoxifen injection to Vil1CreERT2/Mcl1^fl/fl^ mice could represent regeneration from such a pre-existing lineage. Despite efficient recombination taking place in the Vil1CreERT2/Mcl1^fl/fl^ model ((Healy *et al*., 2020) and **Figure 5**), we cannot exclude, however, that it corresponds to proliferation of cells having simply escaped recombination.

Despite an active controversy about the respective roles of quiescent, committed, differentiated or Lgr5-positive cells in intestinal regeneration (Bankaitis *et al*., 2018; Buczacki, 2019), there seems to be a consensus about their common belonging to Lgr5-positive developmental lineage (Ayyaz *et al*., 2019; Nusse *et al*., 2018; Yui *et al*., 2018). Our observation that adult spheroids are not traced by Vil1Cre, nor Lgr5CreERT and that regenerating clones after CBC ablation are similarly not traced by the Vil1Cre transgene, demonstrates that at least a fraction of regenerating crypts originate from an Lgr5-independent lineage.

Identification of cells belonging to this lineage in the homeostatic intestinal epithelium pointed to cells sharing some characteristics with CBCs (Olfm4-positive, abundance of ribosomal protein transcripts) but with downregulation of proliferative markers, suggestive of a quiescent state. Participation of quiescent stem cells to intestinal regeneration has been documented for years (Scoville et al., 2008). They were coined “label retaining cells” from their aptitude to stably incorporate DNA tracers or H2BGFP, and related either to Paneth or enteroendocrine precursors, or to Hopx/Bmi-positive “+4” cells (Buczacki, 2019; Li *et al*., 2016; Roth et al., 2012). Neither Paneth or enteroendocrine cell markers, nor Hopx or Bmi1 transcripts show up in genes differentially expressed in the Vil1Cre-negative cells identified in the present study. This suggests a different origin, in agreement with their Lgr5-independent character. In addition to ribosomal protein genes characteristic of CBCs, the Vil1Cre-negative cluster displays upregulation of nuclear encoded mitochondrial proteins, including complex IV cytochrome oxidases, mitochondrial ATP synthase subunits and the ATPase inhibiting factor. This may sound unexpected, when quiescent stem cells of diverse origin are reported to rely on glycolysis for energy supply (Shyh-Chang and Ng, 2017). However, contrary to what was expected from earlier measurements, quiescent hematopoietic stem cells have been shown to harbor more mitochondria than their more differentiated counterparts, making them ready to rapidly engage in regeneration when needed (Filippi and Ghaffari, 2019; Morganti et al., 2022). An unexpected observation of our study is the effect of expression of the diphtheria toxin receptor (HB-Egf) in CBCs, in the absence of toxin administration. The mere expression of HB-Egf significantly increases the number of cells not traced by the Vil1Cre transgene, suggesting the presence of a chronic injury (Chen et al. 2010). This chronic effect is neglected in the numerous studies using the diphtheria toxin receptor to ablate specifically a cell type, which might deserve re-examination.

In conclusion, our results indicate that a hierarchical stem cell model involving a novel lineage of reserve stem cells applies to regeneration of the intestinal epithelium in addition to the plasticity model.

### Limitations of the study

The fact that the newly identified intestinal lineage is only characterized “by default” (absence of the Vil1Cre transgene expression) constitutes clearly a limitation of our study. While it allowed purification of the corresponding cells from FACS-sorted PlusMinus cells, it makes it difficult to identify these cells in the homeostatic wild-type intestinal epithelium and trace them during development. Future studies exploiting novel lineage-tracing methodologies (Baron and van Oudenaarden, 2019; Jindal et al., 2023; Wagner and Klein, 2020) might allow identification of specific markers of these cells (if such markers exist) which would in turn open the way to their detailed characterization.

## Materials and Methods

### Experimental animals

All the animal procedures complied with the guidelines of the European Union and were approved by the local ethics committee of the faculty of Medicine of Université Libre de Bruxelles (protocols n°713N and n°759N). Mice strains used include: CD1 (Charles Rivers, France); Lgr4/Gpr48^Gt (Leighton et al., 2001); Lgr5-DTR knock in (Tian et al., 2011); Lgr5-CreERT2 (Jackson Laboratory); Lgr5fl/fl (W. et al., 2011); Mcl1fl/fl (Vikstrom et al., 2010b); Tg(Vil1Cre)997Gum/J (Madison et al., 2002); Tg (Vil1Cre/ERT2)23Syr (Sylvie Robine, Paris, France); Rosa26YFP; Rosa26LacZ ; and Rosa26Tomato. 8-16-weeks-old mice were used for these experiments. The day the vaginal plug was observed was considered as embryonic day 0.5 (E0.5).

### Lineage Tracing and specific cell ablation

For lineage tracing experiments, Tamoxifen (Sigma-Aldrich) was dissolved in sunflower oil (Sigma-Aldrich)/ethanol mixture (19:1) at 10mg/ml and used in all experiments at a dose of 0.066 mg/g of body weight. For ablation of CBCs, Lgr5-DTR mice were injected intraperitoneally with 50 µg/kg diphtheria toxin (DT; Sigma-Aldrich). Control mice were either injected with sterile Dulbecco’s phosphate-buffered saline (DPBS, Gibco) solution or not injected.

### *Ex vivo* culture Matrigel domes (old protocol)

For ex vivo organoid culture, the intestine was cut in 3-5 mm pieces and incubated in 1- and 5-mM Ethylenediaminetetraacetic acid (EDTA) in DPBS at 4°C (15 min and 25 min, respectively). Epithelial crypts were cultured according to Sato et al. (Sato et al., 2009) in advanced-DMEM/F12 medium (ThermoFisher scientific) supplemented with 20 mM L-Glutamax (Gibco), 1 X N2 (Gibco), amphotericin, B27 w/0 vit.A (Gibco) and penicillin-streptomycin cocktail, 10 mM N-2-hydroxyethylpiperazine-N-2-ethane sulfonic acid (HEPES), 40 μg/ml gentamycin (all from ThermoFisher scientific), and 1 mM N acetyl cysteine (Sigma). Except if indicated otherwise, the medium contained “ENR” [50 ng/mL EGF, 100 ng/mL Noggin (both from Peprotech), and 100 ng/mL CHO-derived R-spondin1 (R&D System)].

Fetal spheroids were prepared and cultured in ENR-containing medium as described previously (Fernandez Vallone et al. 2016).

Adult spheroids were initially observed when replacing the EDTA step in Sato’s protocol by a Collagenase I (0.132 mg/ml, Sigma-Aldrich)/Dispase (0.66 mg/ml, Gibco) treatment of intestinal tissue for 30 min at 37°C. Thereafter, a two-step protocol was devised as follows: the tissue was processed according to Sato et al (Sato *et al*., 2009), with the EDTA fraction generating organoids; the materials retained on the 70 µm cell strainer (Corning) were washed with Hanks’ Balanced Salt Solution (HBSS, Gibco) and centrifuged at 300 g for 5 min at RT. This washing step was repeated 2 more times. The pellets were treated with 0.132 mg/ml Collagenase, 0.66 mg/ml Dispase, and 0.05 mg/ml DNase1 (Roche Diagnostics Gmbh) dissolved in Dulbecco’s Modified Eagle Medium (DMEM, Gibco) for 15 min at 37°C under agitation at 75 rpm. Mixing by up-and-down pipetting was then performed with a 10 mL pipette before a further 20 min incubation at 37°C. The up and down mixing step was repeated which resulted in complete dissociation. The dissociated samples were passed through a 100 µm cell strainer (VWR) and centrifuged at 300 g for 10 min. The pellets were washed twice with basal culture medium (BCM; containing Advanced-DMEM/F12 medium supplemented with 1 X penicillin-streptomycin cocktail 100X, 1 X amphotericin and 0.8 μl/ml gentamycin) plus 2 mM EDTA (Invitrogen), followed by 2 washes with DPBS (Gibco) containing 10% Fetal Bovine Serum (FBS). All washes were performed at RT and involved centrifugation at 300 g for 5 min. Finally, the resulting fraction was plated in basement membrane matrix, LDEV free Matrigel (Corning). The culture medium for the first 48 h was ENR in advanced-DMEM/F12 medium (Thermo fisher scientific) supplemented with 10 µM Y-27632 (Peprotech), as described above, in all initial seedings. The medium was changed every other day either with ENR or BCM, as reported in the Results section. Pictures were acquired with a Moticam Pro camera connected to a Motic AE31 microscope.

Intestinal fibroblasts were isolated and cultured as described (Powell *et al*., 2011) in ENR medium and their supernatant collected after 3 days. This supernatant was used as conditioned medium. In co-culture experiments, organoids were cultured in medium containing 50% fibroblast conditioned medium with 50% organoid culture medium (V/V, organoid culture medium was made of advanced-DMEM/F12 medium with 2% FBS and ENR).

### Sandwich Matrigel protocol

The protocol was derived from Bues et al. (Bues, Biočanin et al. 2020). The tissue samples were dissociated as described above, the final pellets were dissolved in ENR, and spread in wells previously coated with solidified Matrigel. After 1 h in the incubator (37°C, 5% CO2) the medium with unattached material was removed. A second layer of Matrigel was added on top of the cells attached to the first layer. Following Matrigel solidification, ENR medium supplemented with 10 µM Y-27632 was added to each well. This protocol was used only for initial seeding of adult spheroids.

### Tissue processing and immunohistochemical analysis

The whole small intestine was surgically removed and washed with ice-cold DPBS without Ca+2/Mg+2. The intestine was cut open longitudinally with the mucosa outwards and was rolled around a 1 mL pipette from duodenum toward ileum. The tissue samples were immediately fixed with a 10% formalin solution (VWR), overnight (O/N) at room temperature (RT), and then incubated in succession in 20% and 30% sucrose solution for at least 24 hours each before being embedded in Tissue freezing medium (Leica). Histological and immuno-fluorescence/histochemistry experiments were carried out on 6 μm sections. A 10 mM citrate sodium (pH 6) solution was used as an epitope retrieval solution when required. The tissue samples were incubated with the primary antibodies O/N at 4°C. The secondary anti-species biotin- or fluorochrome-coupled antibodies were incubated for a minimum of 1 h at RT. ABC kit (Vector Labs) and DAB substrate Kit (Vector Labs) were used for target revelation in immunohistochemistry. Hematoxylin (immunohistochemistry) or DAPI (immunofluorescence) were used for nuclei staining according to standard procedures. The primary and secondary antibodies used for staining are listed in **table S5**. After dehydration, slides were mounted in a Xylene-based medium (Coverquick 4000, VWR Chemicals) in case of Immunohistochemistry or with Glycerol Mounting Medium (Dako) containing 2.5% 1,4 Diazabicyclo [2.2.2] octane (Sigma-Aldrich) in case of immunofluorescence. Nanozoomer digital scanner (Hamamatsu) and Zeiss Axio Observer inverted microscope and Aurox spinning disk (immunofluorescence) were used for visualization of the samples. The number of animals used for each experiment is reported in Figures or Figure legends.

Regarding *Ex vivo* culture, two fields for imaging were chosen at the start of the experiment for each animal and pictures were taken at different time points. Settings for organoids and spheroids were always the same (**Fig.4A,C,D,H and G**). X-gal staining was performed for 5h on organoids and spheroids at day6 (**Fig.4B**).

### RNA seq and Gene Set Enrichment Analysis (GSEA)

RNA was extracted using mirVana™ miRNA Isolation Kit (Invitrogen). A total of 400 ng extracted RNA was quality-controlled using Fragment Analyzer 5200 (Agilent technologies). Indexed cDNA libraries were obtained using the TruSeq stranded mRNA LP (Illumina) following manufacturer’s recommendations. The multiplexed libraries were loaded on a NovaSeq 6000 (Illumina) and sequences were produced using a 200 Cycles Kit. Approximately 25 million paired-ends reads were mapped against the mouse reference genome GRCm38 using STAR software to generate read alignments for each sample. Annotations Mus-musculus.GRCm38.90.gtf were acquired from ftp.Ensembl.org. After transcript assembling, gene-level counts were obtained using HTSeq. Genes differentially expressed in adult spheroids (as compared with organoids or fetal spheroids) were identified with EdgeR method (minimum 10 counts per million (CPM), FDR < 0.05), and further analyzed using GSEA MolSig (Broad Institute) (Subramanian et al., 2005) to identify biological processes.

### ATAC-Seq experiment

Adult spheroids and organoids were cultured as described previously. 41000 sorted cells were harvested in 1 mL of PBS + 3% FBS at 4°C. Following centrifugation, cell pellets were resuspended in 100 mL of lysis buffer (Tris HCl 10 mM, NaCl 10 mM, MgCl2 3 mM, Igepal 0.1%) and centrifuged at 500 × g for 25 min at 4°C with the break set at 4. The supernatant was discarded, and nuclei were resuspended in 50 mL of reaction buffer (TDE1 transposase 2.5 mL (Illumina), TDE buffer 25 mL (Illumina)). The reaction proceeded for 30 min at 37°C and was then terminated by the addition of 5 mL of stop buffer (NaCl 900 mM, EDTA 300 mM). DNA purification was performed using the MinElute purification kit (QIAGEN) according to the manufacturer’s instructions. DNA libraries underwent PCR amplification (NEB-Next High-Fidelity 2x PCR Master Mix, New England Biolabs), indexing using previously described primers (Buenrostro et al. 2013) and double-size selection from 150 to 1200 base pairs (bp) using AmpureXP magnetic beads (Beckman) as per the manufacturer’s protocol. The multiplexed libraries were loaded onto a NovaSeq 6000 (Illumina) using a S2 flow cell and paired-end sequences were generated using a 200 Cycle Kit. ATAC-seq paired-end reads were subsequently aligned to the mouse GRCm38 genome using Bowtie2 (version 2.2.6) with specified options. List of ATAC-seq oligos for PCR are listed in **table S5**.

### Gene expression analysis by qPCR and RNAscope

qRT-PCR was performed on total RNA extracted from adult spheroid or organoid cultures using the mirVana™ miRNA Isolation Kit (Invitrogen) according to manufacturer’s recommendations. A DNase I treatment (Invitrogen) was used to remove potential contaminant genomic DNA. cDNA was prepared using RnaseOUT and Superscript II according to manufacturer’s manual (Invitrogen). qTower 3 from Analytik Jena was used to perform qPCR. Gene expression levels were normalized to that of reference genes (Rpl13, Sdha, Ywhaz) and quantified using the qBase Software (Biogazelle). Primer sequences are reported in **table S5**. In situ hybridization experiments were performed based on manufacturer instructions with the RNAscope kit (ACD-Biotechne; probes listed in **table S5**).

### Flow cytometric analysis, cell sorting (FACS) and Single cell RNAseq (scRNAseq)

After removal of EDTA fraction, samples from 5 Vil1Cre (Heterozygotes,HE), Rosa26R-Tomato (HE) mice were treated with collagenase (0.132 mg/ml) Dispase (0.66 mg/ml) and DNase1 (0.05mg/ml) solution for 35 min at 37°C, washed 2 times with BCM-2 mM EDTA solution with centrifugation at 500 g for 2 min in-between. The cell pellet was washed once with PBS-2% FBS (FACS solution) and centrifuged at 500 g for 2 min. The cells were then incubated with rat anti-mouse CD16/CD32 (blocking solution) at a final dilution of 1/50 (v:v) in FACS solution for 15 min. The APC rat anti-mouse CD31 (1/200), APC rat anti-mouse CD45 (1/500) and BV711 rat anti-mouse CD326 (Epcam,1/200) antibodies were added to blocking solution and incubated for 45 min on ice. After 3 washes with FACS solution, DAPI was added to mark the dead cells (1/2000) and the cells were sorted using the FACS Aria I cytometer (BD Biosciences). Twenty thousand Epcam-positive and Tomato-negative (PlusMinus) or Epcam-positive and Tomato-positive (PlusPlus) cells were collected for each animal. These cells were processed through the Chromium Next GEM Single Cell 3’ Reagent Kits v3.1 (10X Genomics) according to manufacturer’s recommendations and sequenced on Novaseq6000 (Illumina). The data were processed through the CellBender software to decrease contamination by ambient RNA (Fleming et al., 2022). Data were analyzed using the Seurat Package in R (Stuart et al., 2019). For PlusMinus sample, 2,650 cells passed the quality control steps (250–2,000 counts and <10% of mitochondrial genes). For PlusPlus sample, 2,154 passed the quality control steps (250–8,000 features and <10% of mitochondrial genes). SCTransform was used as the normalization/scaling method (Hafemeister and Satija, 2019). The UMAPs plots were generated using 15 dimensions and a clustering resolution of 0.4 for the PlusMinus sample, 15 dimensions and a clustering resolution of 0.6 for the PlusPlus sample and 15 dimensions and a clustering of 0.3 for the PlusMinus/PlusPlus merged analysis. Cell types were identified based on previous scRNAseq experiments (Haber *et al*., 2017) and GSEA MolSig (Broad Institute) (Subramanian *et al*., 2005).

### Statistical analyses

Statistical analyses were performed with Graph Pad Prism 9. Venn diagrams in and **Figs.S2,S3 and S6** were performed with Venny, version 2.0.2 (https://bioinfogp.cnb.csic.es/tools/venny/) and analyzed with the hypergeometric p-value calculator (https://systems.crump.ucla.edu/lab-software/). All experimental data are expressed as mean ± SEM. The significance of differences between groups was determined by appropriate parametric or non-parametric tests as described in figure legends.

### Data availability

The datasets generated in this study will be deposited in the Gene Expression Omnibus (GEO) upon publication. They will be accessible under the following accession numbers: GSE262855 and GSE65395 for RNA sequencing data of adult spheroids and organoids, as well as fetal spheroids; GSE262657 for RNA sequencing data of adult spheroids cultured in two-layer Matrigel; GSE262871 for the ATAC-seq experiment; GSE262572 for RNA sequencing data of Lgr5DTR HE and WT; and GSE262537 for single-cell RNA sequencing data.

## Supporting information

Answer to Review Commons reviewers

## Acknowledgements

Special thanks are extended to Christine Dubois for her invaluable assistance with cell sorting experiments, to Fred De Sauvage from Genentech for generously providing access to the Lgr5-DTR mouse line, to Eduardo Andrés Rios Morris for his expertise and support with the Confocal microscope, and also to Andreas Strasser and Philippe Bouillet (The Walter and Eliza Hall Institute of Medical Research, Melbourne, VIC, Australia) for sharing the Mcl1floxed mice and help regarding interpretation of our Mcl1 ISH results. This work was supported in part by grants from the not for profit Association Recherche Biomédicale et Diagnostic (to GV). MM was supported by Fonds pour la formation à la recherche dans l’industrie et dans l’agriculture (FRIA).

## Author contributions

GV conceptualized the study. MM, VFV and GV organized the data, prepared the figures and cowrote the original draft of the manuscript. GaV made the initial observation of Vil1Cre-negative spheroids. MM, VFV and ML performed the experiments. FL and AF performed the bulk and single-cell RNA sequencing (scRNAseq). GD and FL provided guidance for analysis of scRNAseq. AW and JJ provided access to Mcl1floxed mice under the difficult Covid19 conditions. MIG supervised the work of MM and discussed experimental results in real time. All authors reviewed and edited the manuscript.

## Supplementary figures

**Supplementary figure 1.**
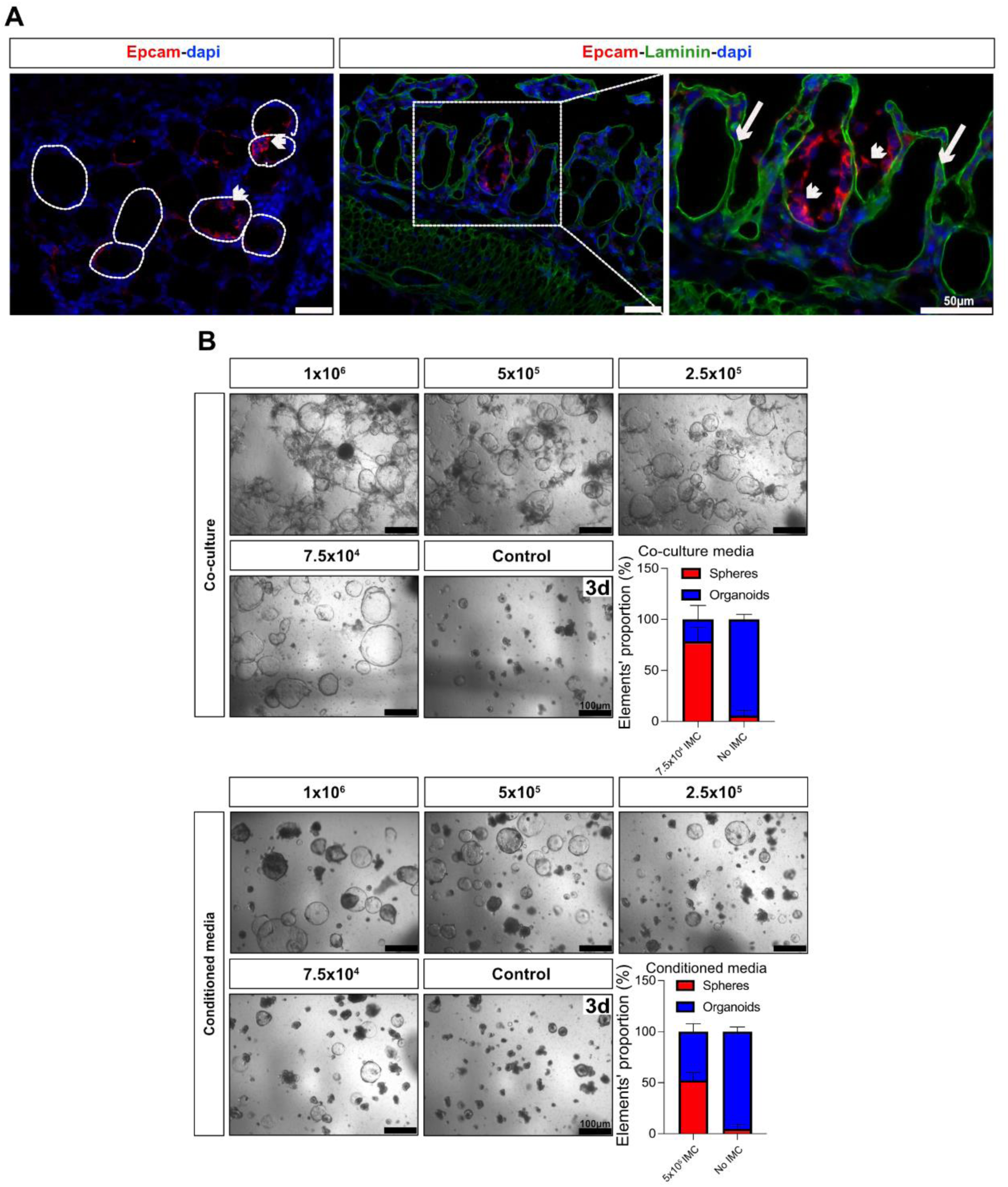
Adult spheroids in ex vivo culture. (A) Immunofluorescence showing section of the material remaining on the filter after EDTA treatment of the intestinal tissue (see figure 1B and Methods); round empty structures correspond to crypt ghosts (dotted contour) surrounded by basal membrane (Laminin) and mesenchyme (arrows), remaining epithelial cells are Epcam positive (arrow heads). (B) Panel showing culture of EDTA-derived organoids in the presence of different numbers of intestinal mesenchymal cells (IMC) in co-culture (upper panel) or in the presence of conditioned media derived from IMC plated at different densities (lower panel); graphs show the proportion of spheroids/organoids elements observed at day3 of culture using 7.5x10^4^ and 5x10^5^ IMC as an example (n=2 different organoids samples for co-culture, n=3 different organoid samples for conditioned media). Scale bars: 50 µm for IF and 100 µm for bright field.

**Supplementary figure 2.**
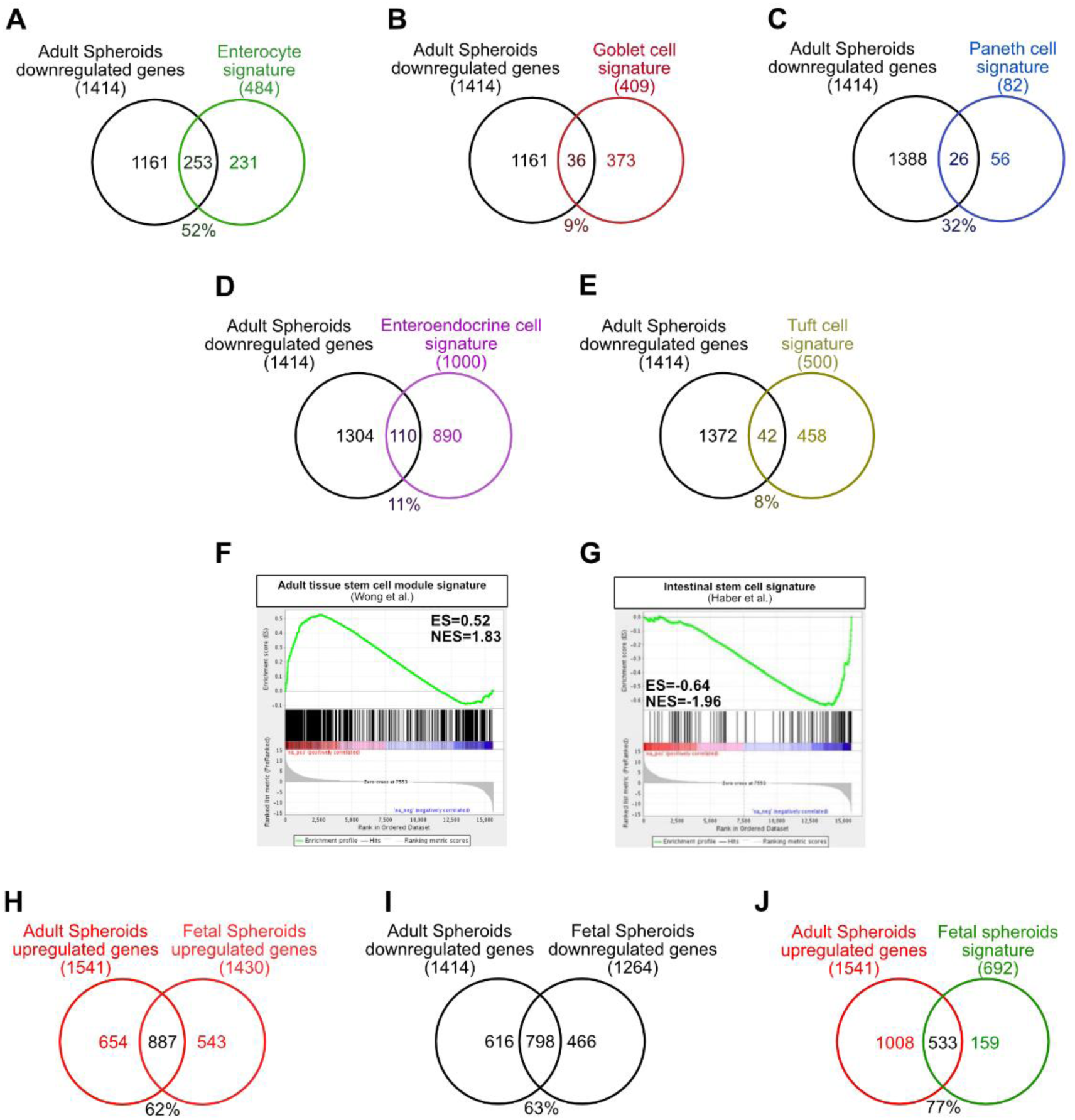
Adult spheroids are made of poorly differentiated intestinal epithelial cells sharing similarities with fetal spheroids. (A-E)Venn diagrams showing the proportion of genes downregulated more than 2-fold in adult spheroids (compared to organoids) that belong to the various differentiated intestinal cell type signatures (Haber *et al*., 2017). Percentages show genes belonging to each signature that are downregulated. (F) Positive correlation of the same set of genes with the adult stem cell module described by Wong et al. (Wong *et al*., 2008) and (G) pre-ranked GSEA analysis showing negative correlation between adult spheroid up-regulated genes (versus organoids) and the intestinal stem cell signature described by Haber et al. (Haber *et al*., 2017). ES = enrichment score, NES = normalized enrichment score. Venn diagrams showing the common number of genes, similarly (H) up- or (I) down-regulated more than 2-fold (when compared to organoids), between adult or fetal intestinal spheroids. (J) Venn diagram showing the number of genes more than 2-fold upregulated in adult intestinal spheroids (when compared to organoids) that belong to the fetal intestinal/stomach spheroid signature previously described (Fernandez *et al*., 2016).

**Supplementary figure 3.**
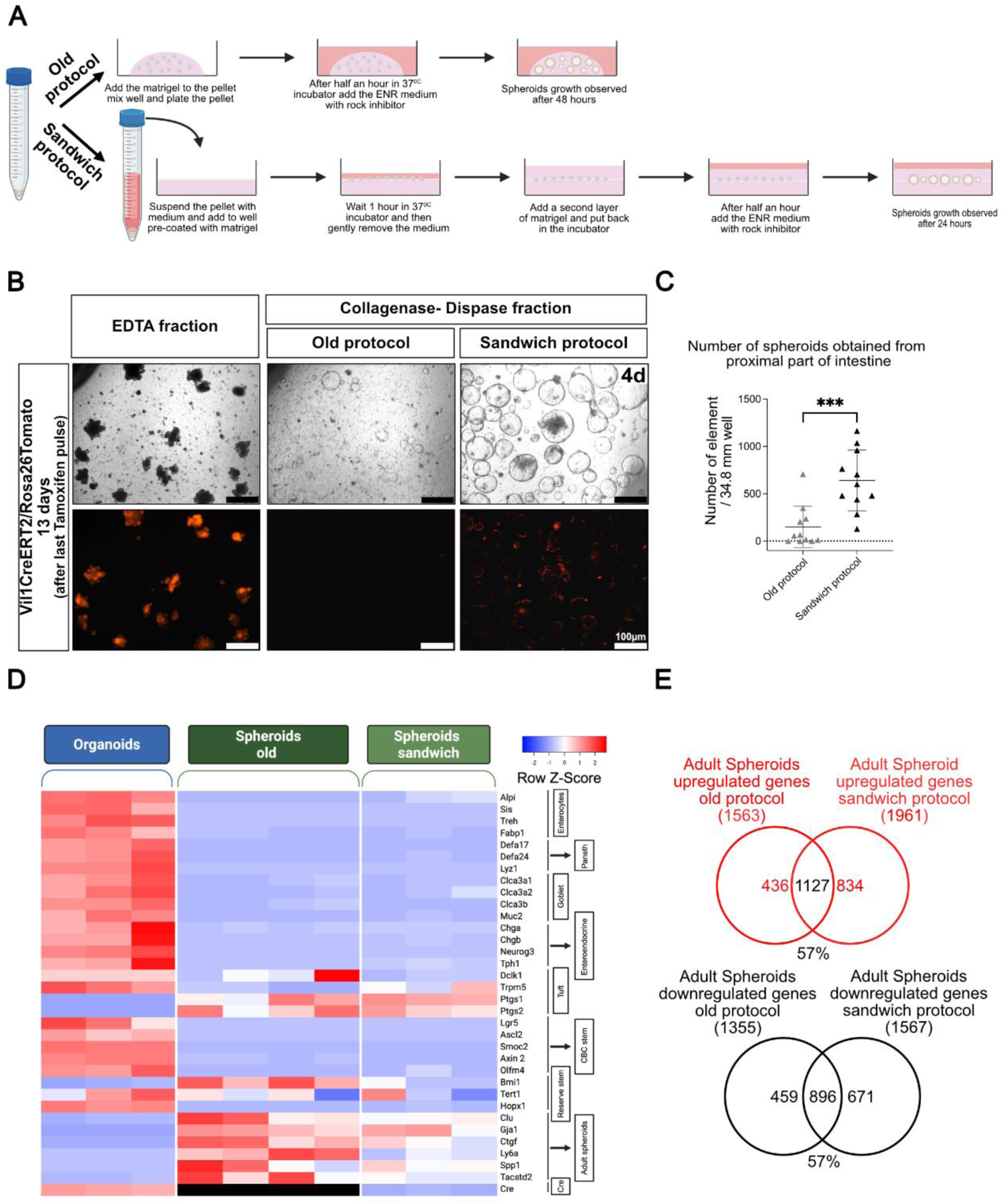
Comparison of adult spheroids obtained with two experimental protocols. (A) Outline of the two protocols. (B) Representative pictures of organoids (EDTA fraction) and spheroids (old or sandwich protocols) of collagenase/dispase fraction obtained from Vil1CreERT2/Rosa26Tomato mice after 3 pulses of Tamoxifen. The animals were sacrificed at day 16 and the pictures were taken at day 4 of culture. (C) Comparison of the yield of spheroids obtained with the two protocols. Each symbol illustrates results from one mouse. (D) Heatmap from bulk RNA sequencing showing similarity between expression of differentiation-, CBC- and spheroid-specific genes in spheroids obtained with the two protocols. Note that Cre transcripts were only assayed in the “new protocol”. (E) Venn diagrams showing the number of genes, similarly up- or down-regulated more than 2-fold when compared to organoids. Statistical analyses: unpaired t test: *** p= 0.0005.

**Supplementary figure 4.**
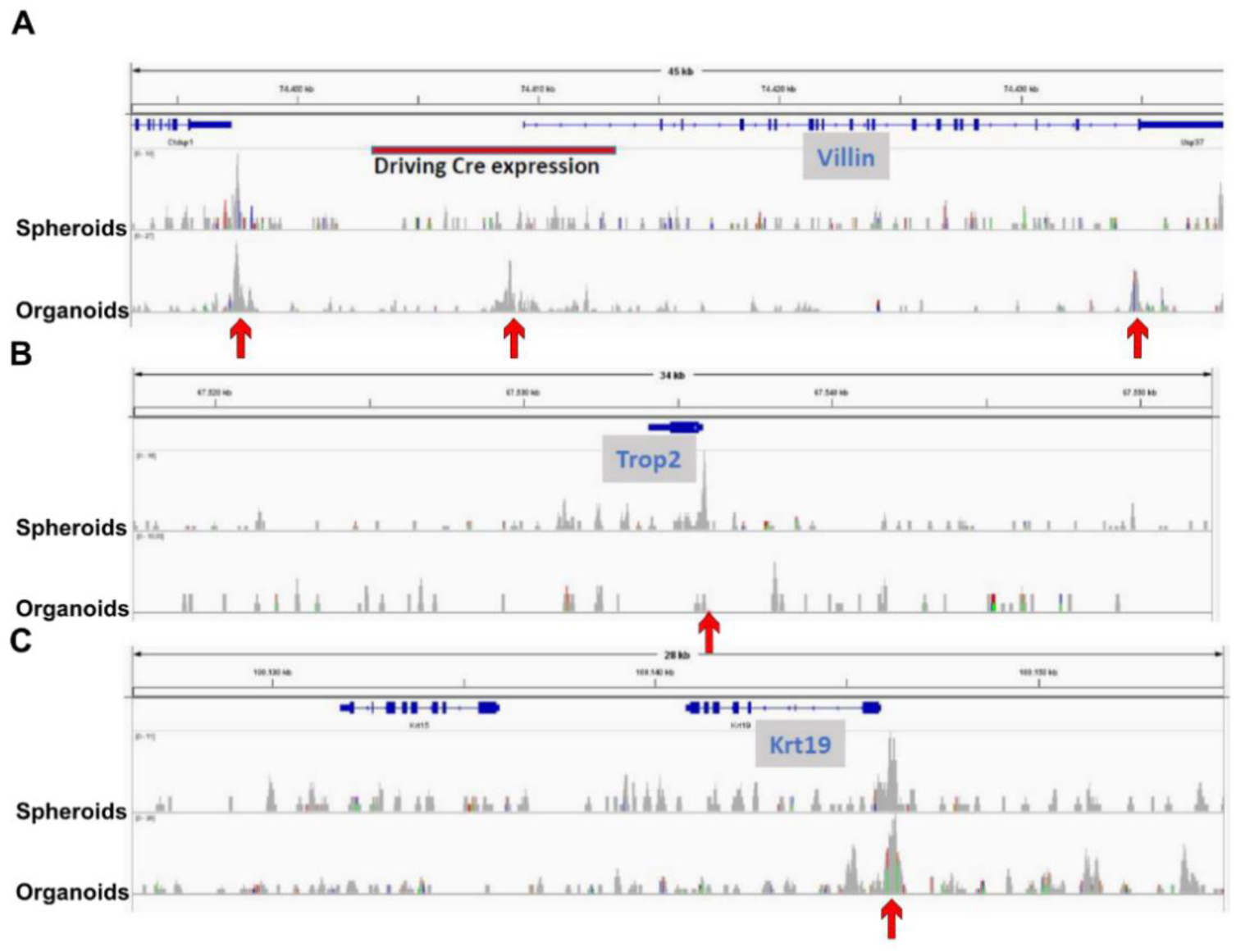
Different organization of chromatin around the villin gene of adult spheroids and organoids. (A) AtacSeq experiment displaying DNA accessibility in the vicinity of the endogenous villin gene from spheroids and organoids in chromatin from a Vil1Cre/Rosa26Tom mouse. (B) Results centered on Trop2 gene and (C) Krt19 gene illustrates control genes, expressed specifically in adult spheroids, or in both spheroids and organoids, respectively. Arrows point to the observed differences.

**Supplementary figure 5.**
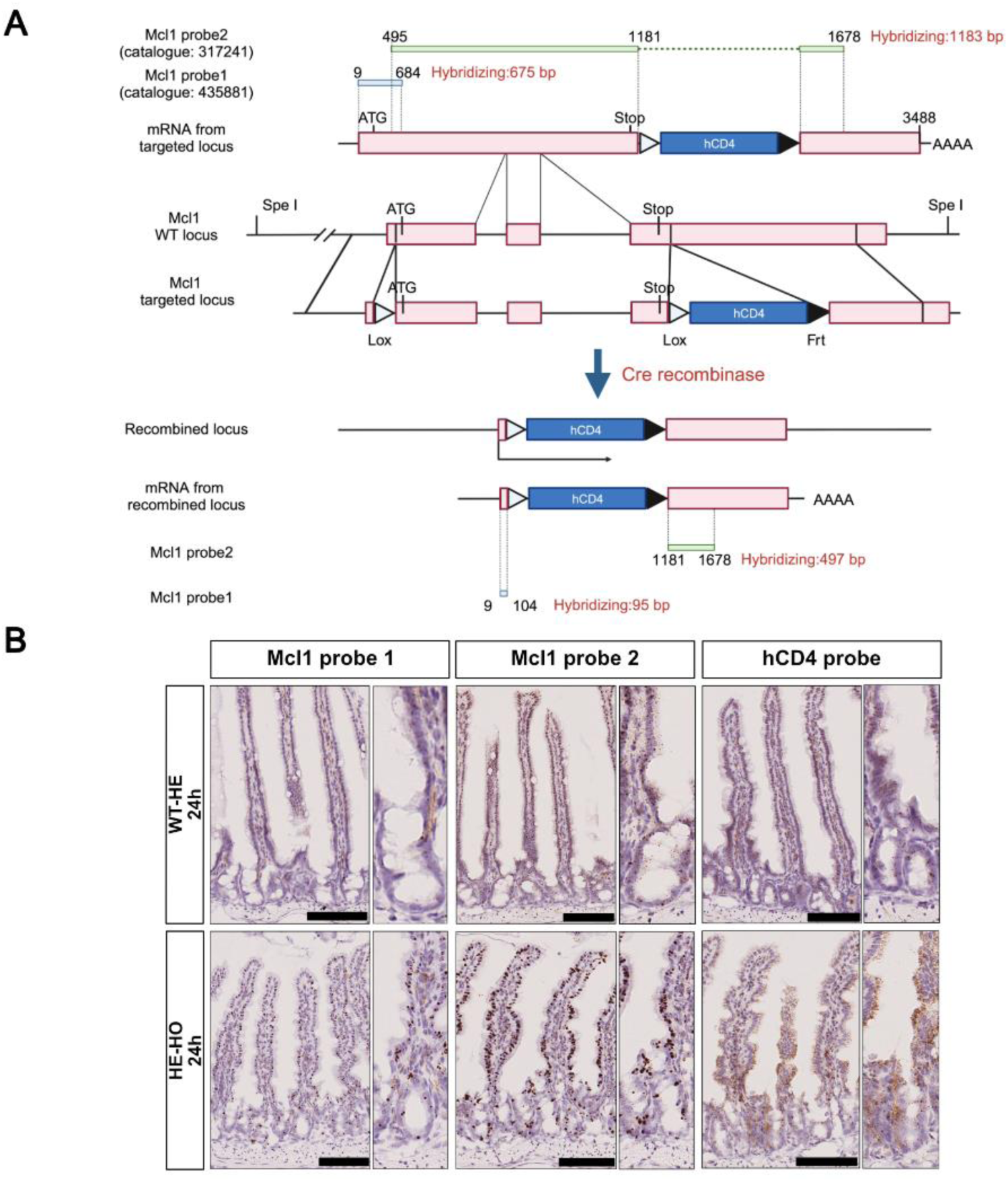
In situ hybridization (ISH) of two Mcl1 probes to mRNA transcribed from non-recombined and recombined Mcl1 loci from Mcl1-floxed mice. (A) Schematic representation of the non-recombined and recombined locus, with the corresponding mRNAs and indication of the extent of the Mcl1 probes. (B) Representative pictures of ISH with the two Mcl1 probes and the hCD4 probe to intestinal sections of Mcl1^fl/wt^ or VilCreERT2/Mcl1^fl/fl^ mice 24 hours after tamoxifen administration, showing the paradoxical increase of the signals.

**Supplementary figure 6.**
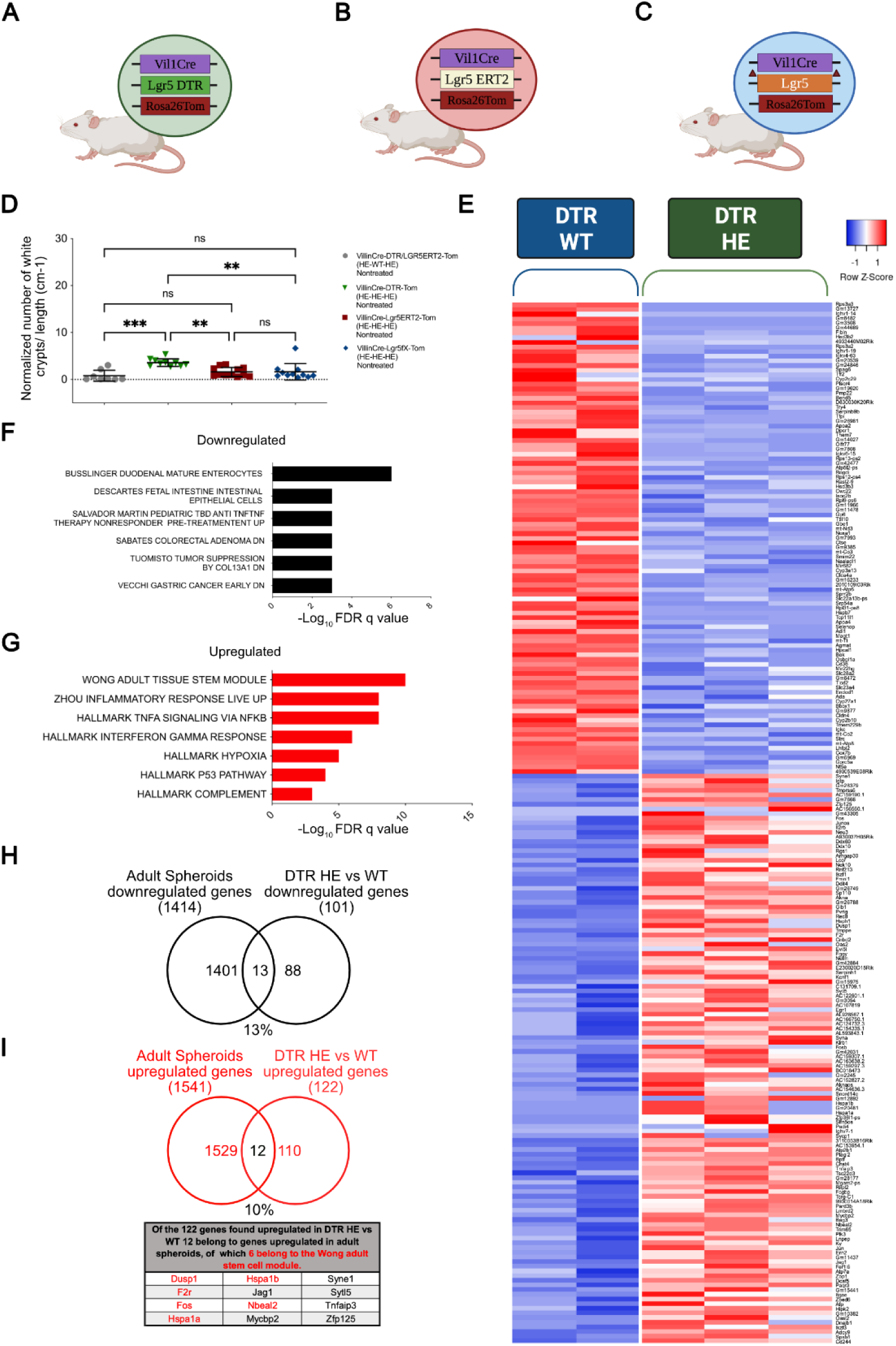
Expression of the diphtheria toxin receptor triggers a regeneration-like phenotype in the intestinal epithelium in the absence of diphtheria toxin. The three transgenic mouse lines used in these experiments were on a Vil1Cre/Rosa26Tom background. All animals were untreated and sacrificed when they were 2 months old. (A) Lgr5-DTR, (B) Lgr5-CreERT2, (C) Lgr5 with a floxed exon 16 allowing its deletion in the Vil1Cre background. (D) Quantification of un-recombined (white) crypts in Vil1Cre/Rosa26Tom double heterozygotes, Vil1Cre/Lgr5DTR/Rosa26Tom, Vil1Cre/Lgr5-CreERT2/Rosa26Tom and VilCre/Lgr5^fl/wt^/Rosa26Tom triple heterozygotes. All mice except VilCre/Rosa26Tom had one Lgr5 allele deleted or non-functional. (E-I) Bulk RNAseq analyses of RNA extracted from uncultured total EDTA-released material; (E) Heatmap from bulk RNAseq of Vil1Cre/Rosa26Tomdouble heterozygotes (DTR WT) compared to Vil1Cre/Lgr5DTR/Rosa26Tom triple heterozygotes (DTR HE). GSEA biological processes of (F) downregulated (C2, C8 cell signature) and (G) upregulated genes (C2 and hallmark genes) were shown in the transcriptome of Lgr5-DTR heterozygous versus wild type crypts. Venn diagrams showing common downregulated (H) or upregulated (I) genes in adult spheroids (versus organoids) and DTR-expressing (versus wild type) intestine. Statistical analyses: One-way ANOVA test with Tukey’s multiple comparison tests: ns p>0.05; *** p=0.0002.

## Supplementary tables

**Table S1.** Bulk RNAseq comparing transcriptomes of adult spheroids and organoids.

**Table S2.** Bulk RNAseq comparing transcriptomes of organoids and adult spheroids cultured in different conditions: ENR, BCM short term (passage 6, day 6), BCM long term (passage 26, day 6).

**Table S3.** Bulk RNAseq comparing transcriptomes of adult and fetal spheroids.

**Table S4.** GSEA MolSig analyses for hallmarks characterizing the adult spheroid transcriptome compared to fetal spheroids.

| Hallmark – Description | *p-value* | FDR *q-value* |
| --- | --- | --- |
| Genes involved in epithelial to mesenchymal transition | 2.46 e<sup>-53</sup> | 1.23 e<sup>-51</sup> |
| Genes regulated by NF-κβ in response to TNFα | 2.45 e<sup>-49</sup> | 6.12 e<sup>-48</sup> |
| Genes up-regulated by KRAS activation | 4.85 e<sup>-32</sup> | 8.09 e<sup>-31</sup> |
| Genes defining inflammatory response | 9.12 e<sup>-30</sup> | 1.14 e<sup>-28</sup> |
| Genes encoding components of blood coagulation system | 8.17 e<sup>-27</sup> | 8.17 e<sup>-26</sup> |
| Genes up-regulated in response to hypoxia | 1.86 e<sup>-26</sup> | 1.33 e<sup>-25</sup> |
| Genes up-regulated in response to IFNγ | 1.86 e<sup>-26</sup> | 1.33 e<sup>-25</sup> |
| Genes up-regulated in response to IFNα proteins | 4.03 e<sup>-23</sup> | 2.52 e<sup>-22</sup> |

**Table S5.**
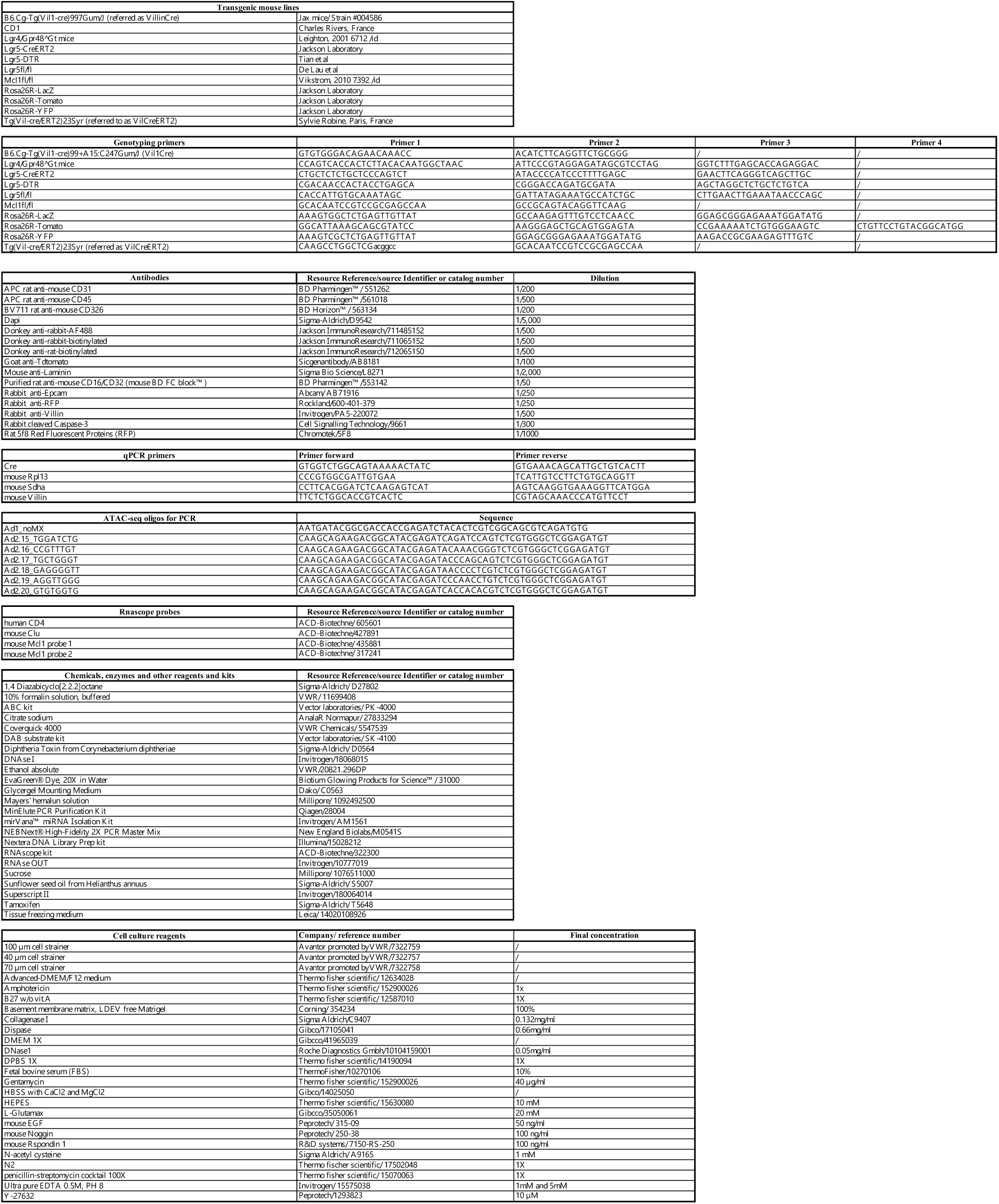
Origin of animals and reagents; list of primers.

