## Supplementary material for "An Lgr5-independent developmental lineage is involved in mouse intestinal regeneration": Answer to Review Commons reviewers

### Revision Plan

**Manuscript number:** RC-2024-02491
**Corresponding author(s):** Gilbert, Vassart

*[The "revision plan" should delineate the revisions that authors intend to carry out in response to the points raised by the referees. It also provides the authors with the opportunity to explain their view of the paper and of the referee reports.*

*The document is important for the editors of affiliate journals when they make a first decision on the transferred manuscript. It will also be useful to readers of the reprint and help them to obtain a balanced view of the paper.*

*If you wish to submit a full revision, please use our "[Full Revision]" template.* ***It is important to use the appropriate template to clearly inform the editors of your intentions.****]*

#### 1. General Statements [optional]

We thank referees 1 and 2 for their in-depth analysis of our manuscript. They see interest in our study, with questions to be answered. Referee 3 is essentially negative, considering that there is nothing new ("*novel finding is missing*").
We respectfully disagree with him/her, comforted by the opinion of referee 2 that "*the authors developed a protocol to reproducibly generate fetal-like spheroids from adult tissue is an important advancement in the field* and … *the manuscript should attract a significant amount of attention in the intestinal field*" and we provide evidence in our answers that he/she did not read the manuscript with the same attention as referees 1 and 2 (see in particular answer to his/her question 5).

Here is a summary of the main reason why we consider that our study represents valuable new information in the field of intestinal regeneration.
It is based on the serendipitous observation that dissociation of adult intestinal tissue by collagenase generates stably replatable spheroids upon culture in matrigel. Surprisingly and contrary to canonical EDTA-generated intestinal organoids and fetal spheroids, these spheroids are not traced in Rosa26Tomato mice harboring a VilCre transgene, despite expressing robustly endogenous Villin. Our interpretation is that adult intestinal spheroids originate from a cell lineage, distinct from the main developmental intestinal lineage, in which the VilCre transgene is unexpectedly not expressed, probaly due to the absence of cis regulatory sequences required for expression in this lineage.
Adult spheroid transcriptome shares a gene signature with the YAP/TAZ signature commonly expressed in models of intestinal regeneration. This led us to look for VilCre negative crypts in the regenerating intestine of Lgr5/DTR mice in which Lgr5-positive stem cells have been ablated

by diphtheria toxin. Numerous VilCre negative clones were observed, identifying a novel lineage of stem cells implicated in intestinal regeneration.
FACS purification and scRNAseq analysis of the rare VilCre negative cells present at homeostasis identified a population of cells with characteristics of quiescent stem cells.

In sum, we believe that our study demonstrates the existence of a hitherto undescribed stem cell lineage involved in intestinal regeneration. It points to the existence of a hierarchical model of intestinal regeneration in addition to the well-accepted plasticity model.

2. Description of the planned revisions

See section 3 below.

3. Description of the revisions that have already been incorporated in the transferred manuscript

*Please insert a point-by-point reply describing the revisions that were already carried out and included in the transferred manuscript. If no revisions have been carried out yet, please leave this section empty.*

Here is a point-by-point reply to the queries of the three referees, with indication of the revisions introduced in the manuscript.

Reviewer #1 (Evidence, reproducibility and clarity (Required)):

In this manuscript, Marefati et al report an Lgr5-independent lineage in the regenerating intestine using in vitro organoids and in vivo injury-coupled lineage tracing model. In organoids, collagenase/dispase dissociated resulted in "immortal spheroids" that maintain a cystic and undifferentiated phenotype in the absence of standard growth factors (Rspondin/Noggin/EGF). Bulk RNAseq of spheroids demonstrates downregulation of classical CBC signatures and upregulation of fetal spheroid, mesenchymal, inflammation and regenerative signatures. In mice, Villin-Cre lineage tracing revealed some Villin- negative progenies that lack reporter tracing throughout crypt-villus ribbons after injury.
The authors proposed that there is Lgr5-independent population support the regenerative response upon CBC depletion. A major caveat of this study is the identification of this population is based on absence of VilCre expression.

We respectfully disagree. It is precisely this characteristic that makes the interest of our study. Whereas mosaicism of transgene expression is widespread and usually of little significance, our study shows that the rare VilCre-negative cells in the intestinal epithelium are not randomly showing this phenotype: they give specifically birth to what we call adult spheroids and regenerating crypts, which cannot be due to chance. The absence of VilCre expression allows tracing these cells from the zygote stage of the various VilCre/Ros26 reporter mice. We have modified our text to emphasize this point.

It is surprising that there is no characterisation of Lgr5 expression throughout the manuscript whilst claiming of a Lgr5- independent lineage.

We understand the perplexity of the referee not to see direct Lgr5 expression data in our manuscript, given our title. However, our point is that it is the cells at the origin of adult spheroids and the regenerating crypts we have identified that are Lgr5-negative, not the spheroids or the regenerated crypts themselves. Those are downstream offspring that may, and indeed have, gained some Lgr5 expression (e.g. figure 3F). We believe that our data showing that VilCre-negative spheroids are not traced in Lgr5-CreERT2/Rosa reporter mice convincingly demonstrate absence of Lgr5 expression in the cells at the origin of adult spheroids (figure 4G). We think that this experiment is better evidence than attempts to show absence of two markers (Tom and Lgr5) in the rare "white" cells present in the epithelium. Regarding the Lgr5 status of cells at the origin of the regenerating "white" crypts that we have identified, the early appearance of these crypts following ablation of CBC (i.e. Lgr5+ve) cells is a strong argument that they originate from Lgr5-negative cells.
Regarding the scRNAseq experiment, Lgr5 transcripts are notoriously low and difficult to measure reliably in CBCs (Haber et al 2017). However, blowing up the pertinent regions of the merged UMAP allows showing some Lgr5 transcripts in clusters 5,6 and none in cluster 1 of figure 8GH. Given the very low level of detection, we had chosen not to include these data in the manuscript, but we hope they may help answer the point of the referee (see portion of UMAP below, with Olfm4 as a control, together with the corresponding violin plot). Several markers that gave significant signals in the CBC cluster (Smoc2, Axin2, Slc12a2) were virtually undetectable in the Olfm4-low /Tom-negative cluster of our scRNAseq data (figure 8I) supporting our conclusion.

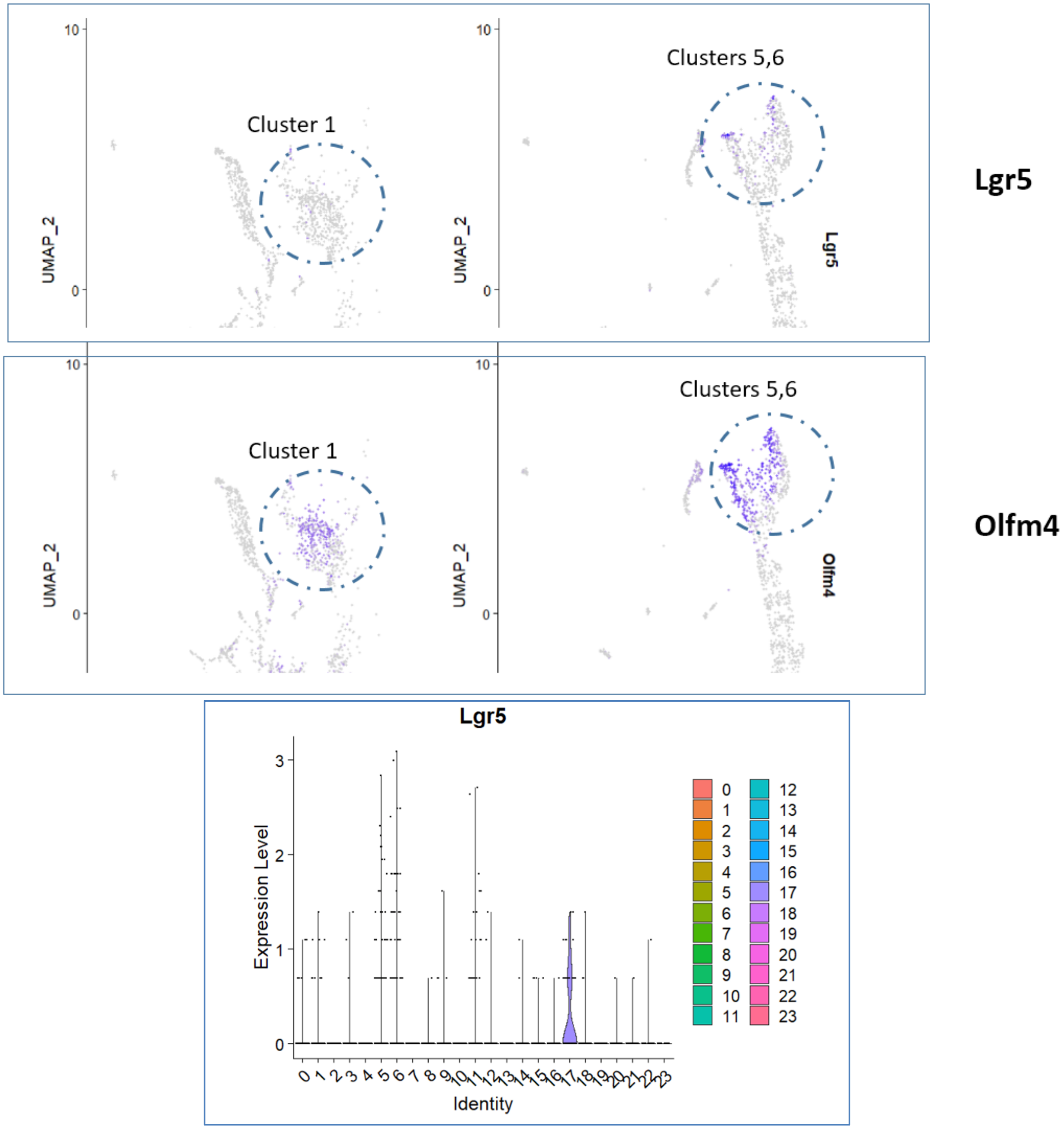

Although the research question is potentially interesting, the concept of epithelial reprogramming upon injury is well documented in the field. The data generated in this manuscript also seem to be preliminary and lack of detailed characterisation. Below are specific comments.

We do not question the existence of epithelial reprogramming upon injury. We believe our data show, in addition to this well demonstrated phenomenon, the existence of rare cells traced by absence of VilCre expression that are at the origin of a developmental cell lineage distinct from Lgr5+ stem cells and also implicated in regeneration.

- Expression of Lgr5 should be properly characterised throughout the manuscript in both organoid models and injury-induced regeneration in vivo.

See above for a detailed answer to this point.

- An important question is the origin of these "Lgr5-independent" adult spheroids. They look and appear like fetal organoids, which could be induced by injury (e.g. upon collagenase/dispase dissociation). Have the authors tried to culture fetal spheroids in BCM over extensive period of time? Do they behave the same? This would be a great way to directly compare the collagenase/dispase-derived organoids with fetal origin.

Fetal spheroids require ENR for survival and die in BCM. We have chosen to illustrate this point in Fig2A by showing that, contrary to adult spheroid, they die even when only Rspondin is missing.

- Fig 1C, Why is the replating spheroid culture time different between mesenchymal cells and conditioned medium?

We took the earliest time showing convincingly the return to the organoid phenotype. This timing difference does not modify the conclusion that EDTA organoids becoming spheroid-like when exposed to factors originating from mesenchymal cells revert to the organoid phenotype when returned to ENR medium without mesenchymal influence.

- It is unclear how the bulk RNA-seq data in Fig. 3 were compared. How long were the adult organoids and spheroids cultured for (how many passages)? Were they culture in the same condition of were they in ENR vs BCM?

Both EDTA organoids and spheroids displaying a stable phenotype were used in this experiment. Organoids were collected at passage 4, day 5; spheroids were collected at passage passage 9 day 3.

As stated in the legend to the figure: "…*to allow pertinent comparison spheroids and organoids were cultured in the same ENR-containing medium…*".

These are important information to consider when interpreting the results. For instance, are Ptgs1 & Ptgs2 expression in adult spheroids the same in ENR vs BCM? Are the gene signatures (regenerative, fetal and YAP) changed in adult spheroids culturing in ENR vs BCM?

We did compare bulk RNAseq of EDTA organoids to ENR-cultured spheroids, short term (passage 6, day 6) BCM-cultured spheroids and long term BCM-cultured (passage 26, day 6) spheroids. To avoid overloading the manuscript these data were not shown in the original manuscript. In summary the BCM-cultured spheroids display a similar phenotype as those cultured in ENR, but with further de-differentiation. See hereafter the results for PTGS, some differentiation markers and fetal regenerative markers including YAP induced genes.

| Symbol | spia log2fold | FDR | orgia-ENR | orgia-ENR | spia-ENR | spia-ENR | spiaBCM-short | spiaBCME-short | spiaBCMLong | spiaBCMLlong |
|---|---|---|---|---|---|---|---|---|---|---|
| Ptgs1 | 8.342183124 | 5.52E-30 | 8 | 5 | 2021 | 2987 | 1634 | 1104 | 3184 | 2603 |
| Ptgs2 | 8.288808842 | 3.06E-11 | 0 | 10 | 895 | 5169 | 1456 | 634 | 959 | 1512 |
| Chga | -4.60963776 | 0.00048245 | 369 | 331 | 64 | 2 | 0 | 14 | 1 | 5 |
| Chgb | -8.99616437 | 2.47E-30 | 2651 | 2297 | 20 | 1 | 4 | 2 | 1 | 1 |
| Alpi | -13.189234 | 7.87E-62 | 8604 | 27628 | 2 | 9 | 0 | 1 | 0 | 0 |
| Muc2 | -5.18847175 | 4.7065E-06 | 758 | 973 | 111 | 2 | 17 | 7 | 4 | 3 |
| Lyz1 | -5.86428178 | 0.00353202 | 1364 | 3703 | 264 | 1 | 1 | 0 | 0 | 0 |
| Lgr5 | -4.03536817 | 0.00506329 | 5766 | 1578 | 667 | 604 | 7 | 4 | 20 | 4 |
| Tacstd2 | 7.714205134 | 1.22E-21 | 8 | 13 | 2377 | 3119 | 3337 | 1328 | 3214 | 883 |
| Ctgf | 7.288713812 | 2.01E-24 | 31 | 88 | 11195 | 10424 | 12651 | 11878 | 6383 | 8418 |
| Legend: | | | | | | | | | | |
| Spia : adult spheroids | | | | | | | | | | |
| Orgia: EDTA organoids | | | | | | | | | | |

We have included a brief description of these data in the new version of the manuscript and added an additional supplementary file (Suppl table 2) presenting the whole data set.

- It is stated: "In agreement with their aptitude to grow indefinitely, adult spheroids express a set of upregulated genes overlapping significantly with an "adult tissue stem cell module" [159/721 genes; q value 2.11 e-94) (Fig.S2F)].". What is the definition of "indefinitely"?
Are they referring to the Fig 1B where spheroid were passaged to P10? The authors should avoid the term "indefinitely" but use a more specific time scale, e.g. passages, months etc.

We agree that the term indefinitely should be avoided, as it is vague. We have introduced the maximum number of passages during which we have maintained the stable spheroid phenotype (26 passages). Also worth noting, the spheroids could be frozen and cultured repeatedly over many months.

SuppFig 3D: Row Z-Score is missing the "e" in Score.

Corrected

- Fig 4E: Figure legend says QNRQ instead of CNRQ.

Corrected

- Fig 4G: The brightfield image of adult spheroids 5 days after 3x TAM injections doesn't look like a spheroid. It seems to be differentiating.

True, the choice was not the best as the spheroids started to darken. When further replated, however, the offspring of these spheroids showing a clear phenotype remain negative 30 days after tamoxifen administration as shown on the figure.

### Revision Plan

We are sorry, but for reasons explained in section 4 below, we cannot redo the experiment to get a better picture.

- Fig 4: Most mouse model data are missing the number of mice & their respective age used for organoid isolation.

We have introduced these data in the legend.

- Fig 4A-D, H-G: How was fluorescent signal of organoids quantified?

The settings of fluo imaging or time of LacZ staining were the same for organoids and spheroid pictures. This has been added to the material and methods of the figure and an example is shown below for Rosa26Tomato.

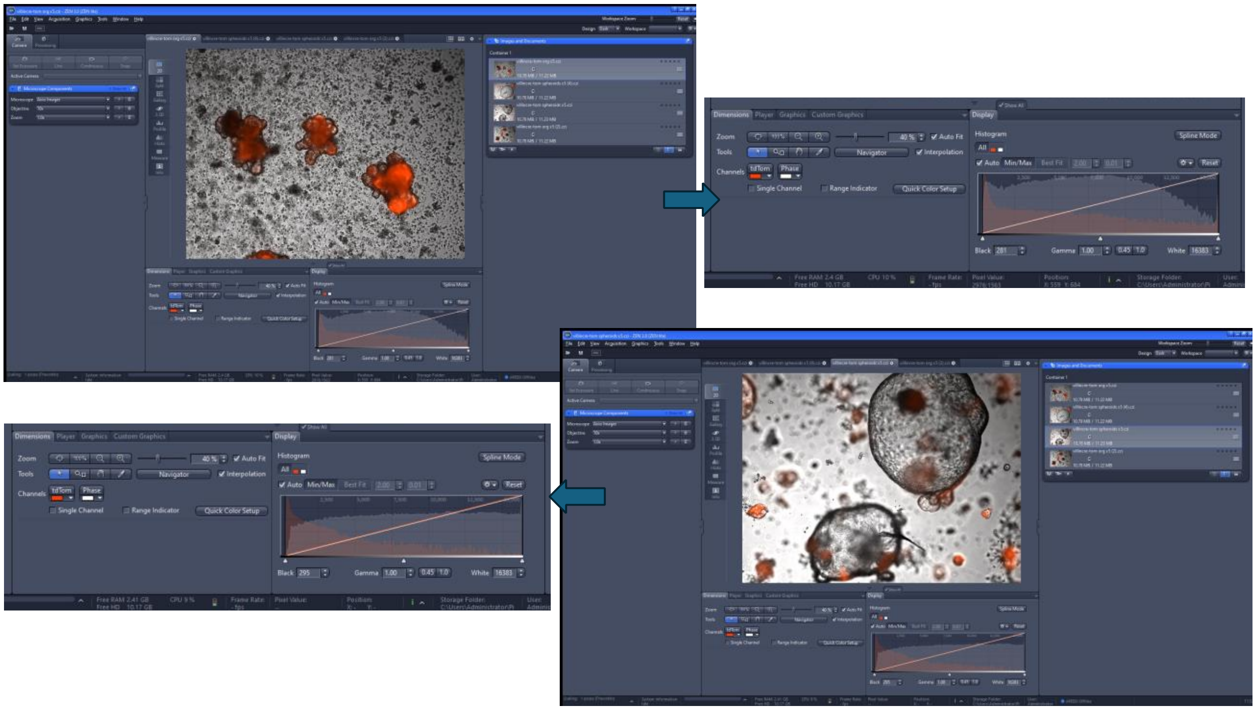

How many images?

2 per animal per condition.

Were there equal numbers of organoids?

No, see number of total elements counted added to the figure

This all needs to be included in methods/figure legends.

We have introduced additional pertinent information in the material and methods section.

- Figure 4B-D, G-H: Which culturing conditions were used for adult spheroids? Original method or sandwich method?

These data were obtained with the original protocol

- Fig 6D-E: Please add the timepoint after DT administration these samples are from. It is not listed in text or figure legend.

These samples were those obtained from mice sacrificed at the end of the 5 day period as indicated in panel A. This has been emphasized in the legend of the figure.

- SuppFig 6D: again timepoint is missing.

In this experiment all samples were untreated as indicated. This has been emphasized in the legend of the figure.

- SuppFig 6: How were the crypts of these mice (DT WT & DT HE) isolated? Was this via EDTA?

This was RNA extracted from total uncultured EDTA-released material (crypts). This has been emphasized in the legend of the figure.

Also, what is the timepoint for isolation for these samples? Even if untreated, the timepoint adds context to the data. Please add more context to describing these different experiments, either in the figure legends or methods section.

All these experiments were from 2 month old animals. We have indicated this in the legend of the figure.

- SuppFig 6E: The quality of the heatmap resolution is too poor to read gene names.

We have improved the resolution of the figure and hope the name of the genes are readable now.

- Fig.5-7, are the regenerating crypt-villus units fully differentiated or are they maintained in the developmental state? Immunostaining of markers for stem cells (Lgr5), differentiated lineages (Alpi, Muc2, Lyz, ChgA etc.) and fetal state (Sca1, Trop2 etc) should be analysed in those "white" unrecombined crypt-villus units.

The differentiation phenotype is shown by the clear presence of morphologically-identified Paneth and Goblet cells. We agree that specific immunostainings could have been performed to further explore this point. Regarding the fetal state, Clu expression was shown during the regeneration period (see figure 7D,E).
Unfortunately, for reasons explained in section 4 below, we are not in a position to perform these additional experiments.

- The following text needs clarification:

"The kinetics of appearance of newly formed un-recombined ("white") crypts was studied after a single pulse of DT (Fig.7A). This demonstrated an increase at 48 hours, with further increase at day 10 and stable maintenance at day 30. The presence of newly formed white crypts one month after toxin administration indicates that the VilCre-negative lineage is developmentally stable and does not turn on the transgene during differentiation of the various epithelial lineages occurring after regeneration (Fig.7B).
Comment: The "newly formed" is an overstatement, the data doesn't conclude that those are "new" crypts.

Except if we do not understand the point, we think we can write that a fraction of “white” crypts must be “newly formed”, since they are in excess of those present in untreated animals at the same time point.

The end of the sentence states that these "white" crypts form developmentally stable lineages, thus these white crypts at day 30 could originate from the initial injury.

As stated above, we consider that crypts found in excess of those present in untreated animals result from the initial injury.

There was no characterisation of the various epitheial lineages. Are they fully differentiated?

See above the point related to Paneth cells and Goblet cells.

Is Lgr5 expressed one month after toxin administration? Can the VilCre neg lineage give rise to CBCs?

We have tried hard to show presence or absence of Lgr5 in white crypts at the various times following DT administration. We tried double RFP / Lgr5-RNA scope labeling and double GFP/RFP immunolabeling. Unfortunately, we could not get these methods to produce convincing specific labeling of CBCs in homeostatic crypts, which explains why we could not reach a conclusion regarding the white crypts.
However, there is an indirect indication that “chronic” white crypts (i.e. those caused by DTR expression in CBC, plus those observed 30 days after DT administration) do not express Lgr5. Indeed, acute regeneration indicated by Clu expression at day 5 in Fig.7C is lower in white crypts than in red ones strongly suggesting that white crypts preexisting DT administration (the “chronic ones) do not express Lgr5DTR.

The relationship between white crypt generation and appearance of Clu-positive revival cells (Ayyaz et al., 2019) was then explored. In agreement with others and similar to what happens in the irradiation model, (Ayyaz et al., 2019; Yuan et al., 2023) Clu-positive cells were rare in crypts of untreated mice and their number transiently increased forty-eight hours after a single pulse of DT, and more so after three pulses of DT (Fig.7C,D).
Comment: Comparing 1 pulse at day 2 vs 3 pulses at day 5 makes the data hard to interpret. How is the Clu ISH level for 1 pulse at day 5? Are they equivalent?

After a single pulse of of DT, Clu is only transiently increased. As shown by Ayyaz et al it is back to the starting point at day 5 (supplementary figure 4 of Ayyaz et al).

Clu-positive cells were less frequently observed in white crypts (see "Total" versus "White" in Fig.7C). This fits with the hypothesis that Clu expression marks acutely regenerating crypts and that a proportion of the white crypts are chronically regenerating due to DTR expression in CBCs."
Comment: I believe the authors suggested that the discrepancy of less Clu expression in white crypts is due to the ectopic expression of DTR in CBCs causing low grade injury without DT administration. This means that some white crypts could have been formed before the administration of DT, and thus are on a different regenerative timeline compared to the white crypts formed from DT administration.

Yes, this is our interpretation. We have clarified it in the text.

Is there any proof of the chronic regeneration? Immunostaining of chronic regenerative markers such as Sca1, Anxa1 or Yap1 nuclear localization would support the claim. It'd be important to show only the white crypts, but not the RFP+ ones, show regenerative markers.

We think that the steady state higher number of white crypts in untreated Lgr5-DTR animals, compared to wild type siblings indicates chronical low-grade regeneration, which is supported by the RNAseq data (Suppl fig6). It must be noted, however, that this phenotype is mild compared to the well described fetal-like regeneration phenotype described in most injury models. Since these white crypts were made at undetermined earlier stages, the great majority of them are not expected to show markers of acute regeneration like Clu, Sca1….

Fig 7D-E: What are the timepoints of harvest for HE-WT-HE 1 pulse DT mice and HE- HE-HE PBS injected mice?

We have added this information in the figure.

- Fig 8-9: Regarding the CBC-like Olfm4 low population, what is the status of Lgr5? This should be shown in the figure since the argument is that this is an Lgr5-independent lineage.

See response to the second point.

And what about the regenerative, Yap, mesenchymal and inflammatory signatures? Are they enriched in the white crypts similar to the in vitro spheroids?

In a portion of white crypts, those we believe are newly formed after CBC ablation (see above), there is a transient increase in Clu, which may be considered a marker of Yap activation. In the CBC-like Olfm4 low cells, as seen by scRNAseq, there is nothing like an actively regenerating phenotype. This is expected, since these cells are coming from homeostatic untreated VilCre/Rosa26Tom animals and are supposed to be quiescent “awaiting to be activated”.

Reviewer #1 (Significance (Required)):

Strengths: The study employed a range of in vitro and in vivo models to test the hypothesis.

Limitations: Unfortunately, the models chosen did not provide sufficient evidence to draw the conclusions. Injury induced reprogramming, both in vivo and in vitro, has been well documented in the field. The new message here is to show that such reprogrammed state is continuous rather than transient; instead of regenerating Lgr5+ stem cells, it can continue to differentiate to all cell lineages in Lgr5-independent manner.

We respectfully disagree with this analysis of our results. What we show is not “that such reprogrammed state is continuous rather than transient; instead of regenerating Lgr5+ stem cells, it can continue to differentiate to all cell lineages in Lgr5-independent manner”, but that a quiescent stem cell line, not previously identified, is activated to regenerate a portion of crypts following CBC ablation.  These cells are not reprogrammed, they correspond to a developmental lineage waiting to be activated and keep their VilCre-negative state at least of 30 days. We believe that their “by default tracing” (VilCre negative from the zygote stage) is as strong an evidence for the existence of such a lineage as positive lineage tracing would be. The increase in crypts originating from this lineage after CBC ablation indicates that it is implicated in regeneration. We do not question the well-demonstrated plasticity-associated reprogramming taking place during regeneration; we simply suggest that this would coexist with the involvement of the quiescent VilCre-negative lineage we have identified.

However, through the manuscript, there was no immunostaining of Lgr5 and other differentiation markers. The conclusion is an overstatement without solid proof.

We have provided the best answer we could to this point in our answer to the second question of the referee hereabove.

Reviewer #2 (Evidence, reproducibility and clarity (Required)):

In this manuscript, the Marefati et al. developed a novel approach to generate spheroids from adult intestinal epithelium using a collagenase/dispase based protocol. Adult spheroids were found to be distinct from classic budding-type organoids normally generated from EDTA based release of the crypt epithelium. Transcriptional profiling indicated that adult spheroids were undifferentiated and similar to regenerating crypts or fetal spheroids. To identify the cell of origin that generates adult spheroids, the authors labelled epithelial cells with VilCreERT-LSL-Tom, VilCre-LSL-GFP and Lgr5CreERT- LSLTom mice. From these experiments the authors conclude that that spheroids are only generated from Vil-Cre negative and Lgr5 negative cells. Next the authors deleted the anti- apoptotic gene Mcl1 using Vil-CreERT mice. This led to a strong apoptotic response throughout the crypt epithelium and tissues processed from knockout mice readily generated spheroids, and in vivo, replenishment of the gut epithelium was mediated by unrecombined cells. In a second model, CBCs were ablated using Lgr5DTR mice and VilCre negative cells were found again to contribute to regeneration of the crypt epithelium. Finally based on the absence of Vil-Cre reporter activity, the authors were able to sort out and perform scRNAseq to profile VilCre negative cells. These cells were found to be quiescent, express the stem cell marker Olfm4 and were also abundant in ribosomal gene expression.

The fact that the authors developed a protocol to reproducibly generate fetal-like spheroids from adult tissue is an important advancement in the field. Previous reports have shown that treatment with various small molecule inhibitors can revert budding organoids into a spheroid morphology, but this manuscript demonstrates that spheroids can also be generated from otherwise untreated cells. This new methodology will provide new tools to dissect the molecular determinants of fetal/regenerative cells in the gut. Based on this, the manuscript should attract a significant amount of attention in the intestinal field.

As pointed out by the authors themselves the study has important limitations that diminish enthusiasm. The primary issue relates to the inability of the team to identify markers of VilCre neg cells other than the fact that these cells are Olfm4+ and quiescent. Nonetheless, for the reasons stated above the manuscript should reach the target audience within the research community, if the authors can address the specific points below related to issues with methodology as well as defining more precisely the characteristics and growth requirements of adult spheroid cultures.

Thank you for this positive analysis of our study.

Major comments

The main conclusion of the study is that Vil-Cre neg cells are rare quiescent Olfm4+ crypt cells. If this is the case, then standard EDTA treatment should release these cells as well. Consequently, spheroids should also emerge from isolated crypts grown in the absence of ENR. If this is not the case how do the authors explain this?

We have tried hard to generate spheroids by culturing EDTA organoids in medium lacking ENR and by treating EDTA organoids with collagenase/dispase, without success. Therefore, we are left with the conclusion that spheroid-generating cells must be more tightly attached to the matrix than those released by EDTA, and that it is their release from this attachment by collagenase that triggers a regeneration-like phenotype. This hypothesis is supported by several models of regeneration in other tissues as indicated in our references (Gilbert et al., 2010; Machado et al., 2021; Montarras et al., 2005).

From the text the authors appear to suggest that growth of adult spheroids is dependent initially on "material" released by collagenase/dispase treatment. An obvious candidate would be mesenchymal cells, which are known to secrete factors such as Wnts and PGE2 that drive spheroid morphology. To test this, the authors should treat spheroid cultures with Porcupine and/or PGE2 inhibitors.

We followed similar reasoning, considering that spheroids express strongly Ptgs1 ,2 (Figure 3A). We thought their phenotype might be maintained by autocrine prostaglandin action. We tested aspirin, a Ptgs inhibitor, which was without effect on the spheroid phenotype. Besides, we explored a wide variety of conditions to test whether they would affect the spheroid phenotype [Aspirin-see above, cAMP agonists/antagonists, YapTaz inhibitors (verteporfin and CA3), valproic acid, Notch inhibitors (DAPT, DBZ, LY511455), all-trans retinoic acid, NFkB inhibitors (TCPA, BMS), TGFbeta inhibitor (SB431542)]. As these results were negative, we did not include them in the manuscript.

If these inhibitors block growth then this would suggest that either stromal cells or autocrine signalling involving these pathways is important. Overall, more in-depth analysis of the growth requirements of adult spheroids is required.

Figure 1d indicates that adult spheroids can be propagated for at least 10 passages. The abstract mentions they are "immortal". The text itself does not address this issue. More precise information as to how long spheroids can be propagated is required. If these cultures can be propagated for 10 passages or more it becomes important to determine what nutrients/mitogens in the basal media are driving growth? Alternatively, what is the evidence that spheroid cultures are completely devoid of mesenchymal cells. The text only mentions that "Upon replating, these spheroids could be stably cultured free of mesenchymal cells (Fig.1B)". No validation is shown to support this.

We agree that “immortal” is not a good way to characterize our spheroids, as also pointed out by referee nr 1. We have changed that in the text, indicating the maximal number of replating we tested was 26 and replacing immortal by stably replatable. Of note, the spheroids could frozen/thawed and recultured many times.
Related to the question whether mesenchymal cells could still contaminate the spheroid cultures, we can provide the following answers:

- No fibroblasts could be seen in replated cultures and multiple spheroids could be repeatedly propagated from a single starting spheroid.
- The bulk RNAseq experiment comparing organoids to ENR or BCM cultured spheroids show, despite expression of several mesenchymal markers (see matrisome in Fig3), absence of significant expression of Pdgfra (see below CP20Millions results from the raw data of new suppl table 2, with Clu, Tacstd2 and Alpi shown as controls).

| Symbol | orgia1-ENR | orgia2-ENR | spia-ENR1 | spia-ENR2 | Spia-BCM-short | Spia-BCM-short | Spia-BCM-long | Spia-BCM-long |
|---|---|---|---|---|---|---|---|---|
| Pdgfra | 3 | 5 | 1 | 3 | 22 | 15 | 11 | 13 |
| Clu | 642 | 2609 | 87154 | 147672 | 427443 | 379002 | 161442 | 172449 |
| Tacstd2 | 8 | 13 | 2377 | 3119 | 3337 | 1328 | 3214 | 883 |
| Alpi | 8604 | 27628 | 2 | 9 | 0 | 1 | 0 | 0 |

- Regarding the nutrients/mitogens in the medium driving spheroid growth, we did not explore the point further than showing that they grow in basal medium (i.e. advanced DMEM), given that the presence of Matrigel makes it difficult to pinpoint what is really needed.

In Figure 2, the authors describe the growth requirements for adult spheroids and indicate that spheroids grown in ENR or EN became dark and shrink. The representative images showing this are clear, but this analysis should be quantified.

Added to the manuscript.

In SF3, the gene expression profile of organoids from the sandwich method only partially overlaps with that of organoids from the old protocol. What are the gene expression differences between the 2 culture systems? Secondly, the sandwich method appears to sustain growth of Tom+ spheroids based on RNAseq and the IF images. This suggest that Vil-Cre negative cells are not necessarily the only source of adult spheroids and thus this experiment seems to indicate that any cell may be converted to grow as a spheroid under the right conditions. These points should be addressed.

Looking back to our data in order to answer the point raised by the referee, we realized that we had inadvertently-compared organoids to ENR-cultured spheroids generated by the first protocol to BCM-cultured spheroids generated by the sandwich method. We have corrected this error in a new version of suppl fig3. This shows increased correspondence between genes up- or downregulated in the spheroids obtained in the two protocols (from 49/48% to 57/57% (Venn diagram on the new figure).
We agree that, even after this correction, the spheroids obtained with the two protocols present sizeable differences in their transcriptome. However, considering the very different way these spheroids were obtained and cultured initially, we do not believe this to be unexpected. The important point in our opinion is that the core of the up- and down-regulated genes typical of the de-differentiation phenotype of adult spheroids is very similar, as shown in the heatmap (which was made with the correct samples!). Also, a key observation is that that both kind of spheroids survive and can be replated in basal medium. As already stated, this characteristic is only seen rare cases [spheroids obtained from rare FACS-purified cells (Smith et al 2018) or helminth-infected intestinal tissue (Nusse et al.2018)]. Together with the observation that the majority of them is not traced by VilCre constitutes what we consider the halmark of the spheroids described in our study.
As shown in figure 4E (old protocol) and Suppl Fig.3 (sandwich protocol) both red and white spheroids were extremely low in VilCre expression. As stated in the text, the fact that some spheroids are nevertheless red is most probably related to the extreme sensitivity of the Rosa26Tom marker to recombination (Liu et al., 2013), but this does not mean that there are two phenotypically different kind of spheroids. It means that the arbitrary threshold of Rosa26Tom recombination introduces an artificial subdivision of spheroids with no phenotypical significance.
Regarding the point made by the referee that "that any cell may be converted to grow as a spheroid under the right conditions", we agree and have shown with others that organoids acquire indeed a spheroid phenotype when cultured for instance in fibroblasts-conditioned medium (see suppl fig1B and (Lahar et al., 2011; Roulis et al., 2020) quoted in the manuscript). However, these spheroids cannot be propagated in basal medium, and revert to an organoid phenotype when put back in ENR (Suppl fig1B).

In Figure 4, the authors conclude that spheroids do not originate from Lgr5 cell derived clones even after 30days post Tam induction. Does this suggest that in vivo and under homeostatic conditions VilCre neg cells are derived from a distinct stem cell pool or are themselves a quiescent stem cell. Given the rarity of VilCre neg cells, the latter seems unlikely.

Despite their rarity, we believe VilCre-negative cells observed under homeostatic conditions are themselves quiescent stem cells. Actually, if they were derived from a larger stem cell pool, this pool should also be VilCre-negative. And we do not see such larger number of VilCre-neg cells under homeostatic conditions.

The problem with the original assertion is that Lgr5-CreERT mice are mosaic and therefore not all Lgr5+ cells are labelled in this model. "White" spheroids may thus derive from cells that in turn derive from these unlabelled Lgr5 cells.

We had considered the possibility that mosaicism [very low for VilCre (Madison et al., 2002); in the 40-50% range for Lgr5CreERT2 (Barker & Clevers. Curr Protoc Stem Cell Biol. 2010 Chapter 5)] could explain our data. We think, however that we can exclude this possibility on the basis that spheroids do not conform to the expected ratio of unrecombined cells, given the observed level of mosaicism. Indeed, for VilCre, a few percent, at most, of unrecombined cells in the epithelium translates into almost 100% unrecombined spheroids. For Lgr5CreERT2 mice, the mosaicism level is in the range of 40%, which is what we observe for EDTA organoids (Figure 4G), while spheroids were in their vast majority unrecombined.
We have included a discussion about the possible role of mosaicism in the new version.

ATACseq experiments were briefly mentioned in the manuscript but unfortunately little information was extracted from this experiment. What does this experiment reveal about the chromatin landscape of adult spheroids relative to normal organoids?

We only performed this experiment to search for an explanation to the paradoxical absence of expression of the VilCre transgene in spheroids, despite robust expression of endogenous villin (Suppl Fig.4). We chose to show the chromatin landscape of a gene equally expressed in both organoids and spheroids (Krt19), a gene specifically expressed in spheroids (Tacstd2) and the endogenous Villin gene also expressed in both. We believe that the observation of a clear difference in pattern of the chromatin accessibility around the endogenous villin gene in organoids and spheroids provides an explanation to the observed results. The cis regulatory sequences needed for expression of the endogenous villin gene seem to be different in organoids and spheroids, which may explain why the regulatory sequences present in the transgene (only 12.4kb) might not allow expression of the transgene in spheroids. We have added a sentence in the manuscript clarifying this point. Missing is obviously the chromatin landscape around the VilCre transgene, but this is beyond reach in such kind of experiments.

Reviewer #2 (Significance (Required)):

The fact that the authors developed a protocol to reproducibly generate fetal-like spheroids from adult tissue is an important advancement in the field. Previous reports have shown that treatment with various small molecule inhibitors can revert budding organoids into a spheroid morphology, but this manuscript demonstrates that spheroids can also be generated from otherwise untreated cells. This new methodology will provide new tools to dissect the molecular determinants of fetal/regenerative cells in the gut. Based on this, the manuscript should attract a significant amount of attention in the intestinal field.

Reviewer #3 (Evidence, reproducibility and clarity (Required)):

### Revision Plan

CR-2024-02491

An Lgr5-independent developmental lineage is involved in mouse intestinal regeneration
Marefati et al.
Homeostatic maintenance of the intestinal epithelium has long been thought to rely upon Wnt signaling responsive Lgr5-expressing stem cells that reside at the crypt base. However, myriad reported mechanisms or populations have been reported to underlie epithelial regeneration after injury. Many groups have reported that reacquisition of a fetal- link intestinal phenotype is an import part of the regenerative response, however the originating cell type has not been definitively identified. Herein, the authors demonstrate that cells from adult homeostatic intestine can generate immortal spheroids that resemble fetal spheroids and are derived independent of Lgr5+ intestinal stem cells (ISCs). The authors then draw the conclusion that this indicates that a hierarchical stem cell model applies to regeneration of the intestinal epithelium, in addition to the plasticity model.

Comments:
1. Please indicate what species is used for studies in Fig 1.

All experiments were performed in Mus musculus.

Please clarify if Figure 2 studies utilize Matrigel or not.

Yes

2. RNA-seq analyses of adult intestinal generated spheroids lack the granularity of single cell analyses and thus it is unclear if this is a homogeneous population or if the population has diversity across it (i.e., enteroids/organoids have a high level of diversity). Many of the conclusions from the RNA-seq study are broad and generalized-for example Fig 3F indicates that markers of the +4 ISC populations (Bmi1, tert, lrig1, hopx) were all expressed similarly in adult spheroids as compared to adult organoids. However, while this may be true in the bulk-RNA-seq analyses, clearly scRNA-seq would provide a better foundation to make this statement, as enteroids/organoids are comprised of heterogeneous subpopulations. . .and it might indicate that these +4 markers have only very low expression in the spheroids. Based upon these concerns, misconclusions are likely to be drawn.

We agree and it would be certainly worthwhile to perform scRNAseq of adult spheroid populations. This would certainly be worth doing in future studies to explore the possible heterogeneity of adult spheroids. We nevertheless believe that our scRNAseq performed on homeostatic intestinal tissue from VilCre/Rosa26Tom mice identify Olfm4-low VilCre-neg cells that are likely at the origin of adult spheroids and display a quite homogenous phenotype.

3. The language around Figure 4 results is confusing. Please define "white" and "red". It might be simpler to designate recombined versus not recombined lineage.

We have clarified this in the figure.

4. The hypothesis that collagenase/dispase solution acts as a proxy for injury is not demonstrated and backed by data. Thus, it is difficult to make the conclusion that this approach could represent a "stable avatar" of intestinal regenerating cells. It is clear that subpopulations of crypt-based cells generate spheroids in culture without collagenase/dispase (see the cited reference Smith et al, 2018).

Smith et al demonstrate clearly the possibility to obtain spheroids with properties probably similar to ours from EDTA derived intestinal crypt cells. However they need to prepurify them by FACS. Besides, Nusse et al describe spheroids similar to ours after infection of the intestine by helminths (Nusse et al. 2018). In our case, and for most labs preparing enteroids with the EDTA protocol, the result is close to 100% organoids. Even if we treat EDTA organoids with collagenase, we do not obtain spheroids. This brought us to the conclusion that spheroid-generating cells must be more tightly attached to the matrix than CBCs and that it is their release from the matrix that activates the spheroid regeneration-like phenotype. This hypothesis is supported by several models of regeneration in other tissues as indicated in our references (Gilbert et al., 2010; Machado et al., 2021; Montarras et al., 2005)

5. A study based on the absence of recombination in a VilCre lineage tracing scenario is not well-established to be strong experimental approach, as there are many reasons why recombination may not cells may not be lineage marked. In order to use this system as the authors intend, they first need to demonstrate that villin is not expressed in the discrete cell population that they are targeting. For the presented observational studies, this would be difficult to do. While they do demonstrate differences in chromatin accessibility between cells from organoids versus spheroids (fig s4), some of these differences could merely be due to the bulk analytical nature of the study and the lack of comparing stem cell populations from spheroids to stem cell populations from organoids-since the spheroids are likely homogenous versus the organoids that only have a small fraction of stem cells-and thus represent a mix of stem cell and differentiated cell populations. The authors do not demonstrate that villin protein expression varies in these cells.
If it were found that villin is not expressed in their "novel" population, then one would expect that the downstream use of villin-based recombination would demonstrate the same recombination potential (i.e., Mcl1 would not be recombined). Both recombination studies in Fig 6 are difficult to interpret, and thus it is not clear if these studies support the stated conclusions. Quantification of number of crypts that are negative should be reported as a percentage of recombined crypts.

We are sorry but there seems to be a complete misunderstanding of our data regarding the point raised by the referee. The important point of our initial observation is that *despite robust expression of villin in spheroids, the VilCre transgene is not expressed* (see figure 4E). This in our opinion makes absence of VilCre expression (or of Rosa marker recombination) a trustful marker of a new developmental lineage. All the data in figure 4 constitute an answer.

The reasoning about heterogeneity of cell type in organoids versus probable homogeneity of spheroids is well taken. However, as the endogenous villin gene is expressed in all cells of both organoids and spheroids, it is highly significant that only spheroids do not express the

transgene.
We performed the ATACseq experiment to search for an explanation to the paradoxical absence of expression of the VilCre transgene in spheroids, despite robust expression of endogenous villin (Suppl Fig.4). We chose to show the chromatin landscape of a gene equally expressed in both organoids and spheroids (Krt19), a gene specifically expressed in spheroids (Tacstd2) and the endogenous Villin gene also expressed in both. We believe that the observation of a clear difference in pattern of the chromatin accessibility around the endogenous villin gene in organoids and spheroids provides an explanation to the observed results. The cis regulatory sequences needed for expression of the endogenous villin gene seem to be different in organoids and spheroids, which may explain why the regulatory sequences present in the transgene (only 12.4kb) might not allow expression of the transgene in spheroids. We have added a sentence in the manuscript clarifying this point. Missing is obviously the chromatin landscape around the VilCre transgene, but this is beyond reach in such kind of experiments.

6. Figure 8 indicates that the cell population identified by scRNA-seq may be quiescent. Companion IF or IHC should be conducted to confirm this finding, as well as other conclusions from the informatics conducted.

We agree that additional experiments could be performed to support this point. We are unfortunately not in a position to perform these experiments (see section 4 below).

7. Clearly the data is intriguing, however, the conclusion is strong and is an over interpretation of the presented data. There are a number of validation or extension data that would enhance the overall interpretation of the study:
a. validation of scRNA-seq or bulk RNA-seq concepts by protein staining of intestinal tissues in the damage model will serve as a secondary observation.
b. identification of the ISC that they are defining is critical and important. There is already the notion that this cell type exists and it has been shown with various different markers.
c. expand the analyses of the fetal-like expression profiling to injured intestines to demonstrate that the lineage negative cells indeed express fetal-like proteins.
d. expand the discussion of the Clu+ cell type. Is this cell the previously described revival cell? If so, how does this body of work provide unique aspects to the field?

We agree that all these suggested experiments could be performed and would be of interest. However, we consider that they would not modify the main message of our study and would only constitute an expansion of the present work. As already stated, we are not in the position to perform them (see section 4).

8. There is some level of conflicting data, with the stem population being proliferative in culture stimulated by the stromal cells, but quiescent in vivo and also based upon scRNA-seq data in Fig 9.

We do not see any conflict in our observation regarding this point. The observation that cells that are quiescent in vivo become proliferative when subjected to culture (with or without addition of stromal cells) is routinely made in a multitude of cell culture systems. In particular, it has been shown that intestinal tissue dissociation activates the Yap/Taz pathway, resulting in proliferation (Yu et al. Hippo Pathway Regulation of Gastrointestinal Tissues. Annual Review of Physiology, 2015 Volume 77, 201-227).

9. Many of the findings have been previously reported: Population that grows as spheroids (Figure 2), Population that is Wnt independent (Figure 2), Lgr5 independent regenerative growth of the intestine (figure 3F, Figure 4), Clu+ ISCs drive regeneration (Figure 7).

Whereas these individual findings have indeed been reported, it was in a different context. We strongly disagree with the underlying suggestion that our study would not bring new information. We have identified here a developmental lineage involved in intestinal regeneration that has not been described up to now.

Minor comments:
1. The statement that spheroids must originate from collagenase/dispase digested material might be an overstatement. As spheroids generation from EDTA treated intestines have been previously reported (Smith et al, 2018).

See answer to point 4 above.

Overall while the study includes an extensive amount of work and different approaches, a foundationally supported novel finding is missing. Many of the statements have already been demonstrated by others in the fields. In addition, one of the most intriguing aspects of the study is that the stromal population impacts this stem cell population, however, interactions and factors stimulating the crosstalk are not addressed.

Reviewer #3 (Significance (Required)):

Overall while the study includes an extensive amount of work and different approaches, a foundationally supported novel finding is missing. Many of the statements have already been demonstrated by others in the fields. In addition, one of the most intriguing aspects of the study is that the stromal population impacts this stem cell population, however, interactions and factors stimulating the crosstalk are not addressed.

We can only disagree.

#### 4. Description of analyses that authors prefer not to carry out

*Please include a point-by-point response explaining why some of the requested data or additional analyses <u>might not be necessary or cannot be provided within the scope of a revision</u>. This can be due to time or resource limitations or in case of disagreement about the necessity of such additional data given the scope of the study. Please leave empty if not applicable.*

### Revision Plan

We have answered most questions raised by the referees by explaining our view, by clarifying individual points and, in several cases, by providing additional information that was not included in the original manuscript.
In a limited number of cases when additional experiments were suggested, we were unfortunately obliged to write that we are not in a position to perform them. This is because my lab is closing after more than fifty years of uninterrupted activity. There will unfortunately be nobody to perform additional experiments.
Nevertheless, as written by referees 1 and 2, we believe that the revised manuscript, as it stands, contains data that will be of interest to the people in the field and may be the bases for future developments. We hope editors will find interest in publishing it.
